# Structural and functional landscape of α-synuclein fibril conformations amplified from cerebrospinal fluid

**DOI:** 10.1101/2022.07.13.499896

**Authors:** Arpine Sokratian, Ye Zhou, Enquan Xu, Elizabeth Viverette, Lucas Dillard, Yuan Yuan, Joshua Y. Li, Ariana Matarangas, Jonathan Bouvette, Mario Borgnia, Alberto Bartesaghi, Andrew West

**Author notes:** These authors contributed equally to this study (Arpine Sokratian, Ye Zhou).

## Abstract

Lewy body dementias are pathologically defined by the deposition of α-synuclein fibrils into inclusions throughout the brain. Cerebrospinal fluid(CSF) in disease harbors circulating α-synuclein-fibril seeds, and parental α-synuclein fibrils can template core structure into amplified fibrils. Using cryo-electron microscopy, we identify six novel α-synuclein fibril assemblies amplified from ten CSF samples (3.8Å to 2.9Å nominal resolutions). Fibrils are classified based on two types of filament interaction, two types of β-sheet stacking, and two types of hydrophobic pocket. CSF-amplified fibril products have one, two, or three distinct assemblies each. Six of ten samples share a common fibril assembly. Within this classification, the fibrils have distinct profiles in amyloid dye binding, and dramatically different potencies in both seeding new inclusions in neurons and evoked microglial pro-inflammatory responses. However, no single structural feature predicts functional phenotypes. Our results highlight CSF as a valuable resource to identify novel α-synuclein assemblies potentially important in disease.

## Introduction

α-Synuclein is a 140-amino-acid protein localized to presynaptic nerve terminals in the brain and involved in vesicle and membrane binding^1^. Structurally, α-synuclein adopts amphipathic α-helical loops as well as disordered soluble conformations in the cytoplasm^2,3^. In Lewy body diseases including Parkinson’s disease (PD), dementia with Lewy bodies (DLB), and Parkinson’s disease with Dementia (PDD), soluble α-synuclein undergoes conformational changes that can include the formation of stable antiparallel β-sheet fibrils^4^. As with other proteins that can fibrilize, initial formation occurs through a nucleation-dependent polymerization reaction^5–7^. α-Synuclein fibrils spread inside cells and from cell to cell in a prion-like manner, recruiting endogenous low molecular weight α-synuclein (e.g., monomer) into new α-synuclein fibrils^8,9^. Myeloid cells of the immune system (e.g., microglia) elicit strong pro-inflammatory responses to low concentrations of α-synuclein fibrils that are also thought to contribute to neurodegenerative disease processes. In contrast, other α-synuclein conformations (e.g., monomer, oligomeric or larger truncated species) are comparatively benign with respect to immunogenicity in model systems^10^.

Emerging studies in different models suggest that α-synuclein fibrils grown in the presence of existing or pre-formed fibrils do not have random structural characteristics but instead inherit the core structural features associated with the parental fibrils^11–14^. This property is not unique to α-synuclein fibrils since β-Amyloid and insulin protein fibrils also faithfully transfer distinct structural characteristics to progeny fibrils, even after many rounds of template-nucleated amplification *in vitro* or in cells^5–7,15,16^. With tau protein, *in vitro* amplification from hyperphosphorylated 2N4R-tau isoform seeds conserves disease-specific bioactive characteristics without requirement of post-translationally modified protein in new fibril construction^17^. α-Synuclein fibrils grown from E46K-mutated α-synuclein imprint the aberrant E46K-fibril structure into new fibrils composed exclusively of WT-α-synuclein^12^, suggesting that fibril structural differences caused by an amino acid sequence variant may imprint into new fibrils that lack the sequence variant from the starting seed assemblies. In ground-breaking work, α-synuclein fibrils amplified from brain tissue homogenates from Lewy body diseases are diverse and different from multiple-system atrophy (MSA) according to fluorescent probe binding, NMR spectroscopy, and electron paramagnetic resonance analysis^18^. One study suggests fibril strains purified from MSA are ~1,000 fold more potent in seeding activity in model systems than strains purified from Lewy body diseases^19^.

The spectrum of functional properties encoded within α-synuclein fibrils associated with Lewy body diseases are not well understood, but further insights might reveal new assemblies that may affect disease phenotypes. It has been suggested that lower functional potency associated with PD and DLB fibril assemblies compared to MSA assemblies makes them more difficult to study^20^. However, particularly with DLB and PDD, very high α-synuclein fibril-seeding activity is observed in cerebrospinal fluid (CSF) in different types of *in vitro* aggregation seeding assays. In contrast, healthy controls without neurodegenerative diseases lack detectable α-synuclein seeding activity in CSF^21–23^. Importantly, α-Synuclein fibril seeding activity in CSF from PD cases can be mitigated with the prior incubation of anti-α-synuclein antibodies^24^, suggesting the presence of circulating α-synuclein fibril seeds in CSF. Interestingly, *in vitro* seeding activity in CSF from MSA cases is weak or absent^25,26^, and *in vitro* amplified MSA fibril strains amplified from tissue-procured MSA fibril seeds do not appear to resemble the parental twisted filaments well^27^. Thus, the presence of α-synuclein fibril seeding activity in CSF appears to differentiate Lewy body diseases from MSA, with high specificity and sensitivity both early in disease and throughout the disease course^24,28^.

In this study, we harness the high α-synuclein fibril seeding activity inherent to CSF from pathologically confirmed Lewy body dementia cases to survey the structural landscape of novel α-synuclein fibril assemblies using cryo-electron microscopy analysis coupled with quantitative functional assays in primary cells. With this approach, we identify six novel α-synuclein fibril assemblies amplified from ten subjects, typically present with other fibril assemblies identified in past reports. The novel α-synuclein fibril assemblies have strikingly different potencies with respect to seeding activity in neurons as well as eliciting pro-inflammatory responses in human microglia. However, no single structural feature we could identify fully predicts the functional responses, highlighting the inherently diverse landscape of α-synuclein fibril assemblies in Lewy body dementia. Our results highlight CSF as a potentially valuable resource to study patientspecific fibril compositions and how they might change over time in the course of disease.

## Results

### α-Synuclein fibrils amplified from DLB-CSF segregate into two distinct fibril classes

We selected ten DLB-CSF samples previously ranked for α-synuclein fibril seeding activity in fibrilization assays from pathologically confirmed neocortical-type DLB cases with a high burden of Lewy pathology across the brain (**Table 1** and^23^). Although the CSF samples are postmortem, comparable (age, sex, PMI, etc.) post-mortem CSF from controls without neurological disease had no α-synuclein fibril seeding activity^23^. Combining high-purity and low-endotoxin human WT-α-synuclein monomer protein (**Supplemental Fig. 1**) with DLB-CSF (**Fig. 1a**) in phosphate-buffered saline (PBS), “DLB-I” has the highest seeding, followed by intermediate activities in “DLB-II through V” cases, and lower seeding activities with “DLB VI-X” (**Fig. 1b**). Incubations in the same assay with CSF from a neurologically normal control subject does not yield fibril product in 60 hours, ensuring that the resultant α-synuclein fibril products amplified from DLB-CSF are not the result of spontaneous nucleation. Fibril products amplified from DLB-CSF without thioflavin-T dye show dominant twisted filaments amenable to cryo-EM structural analysis **(Fig. 1c**).

**Figure 1.**
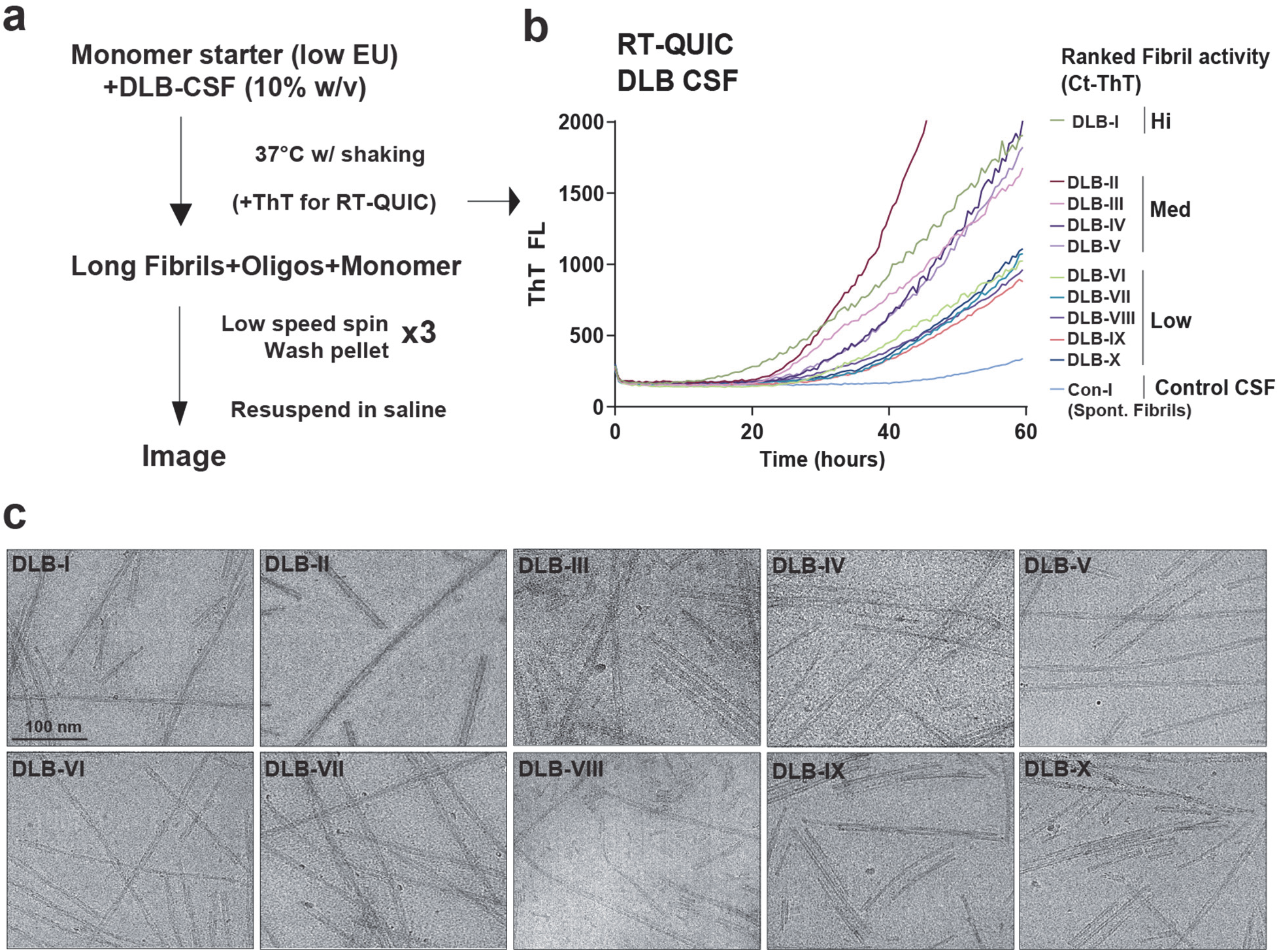
Characterization of α-synuclein fibril products amplified with CSF from Lewy body dementia cases. CSF samples were obtained post-mortem from pathologically defined neocortical-type dementia with Lewy bodies (DLB) cases, or a neuropathologically normal and clinically neurologically normal control known to lack α-synuclein seeding activity (see Table 1). **a.** Procedure for the amplification of α-synuclein fibrils, where monomeric α-synuclein (see **Supplemental Fig. 1**) is combined with 10% (w/v) post-mortem CSF and incubated at 37°C under continuous shaking for 60 hours. Heavy fibril assemblies formed during this time are pelleted with low-speed centrifugation and washed three times in saline. **b.** Representative incubations monitored with thioflavin-T fluorescence from CSF samples previously identified with high, intermediate, or lower α-synuclein seeding activity ^23^. **c.** Representative transmission electron micrographs of α-synuclein fibril products amplified from CSF without thioflavin-T dye included in the incubation (scale bar is 100 nm).

To assess the structural topology of the DLB-CSF amplified α-synuclein fibrils, we performed cryo-EM analysis on single large batches of amplified products from DBL-CSF I-X samples. A recent cryo-EM study by Guerrero-Ferreira et al^29^ examining *in vitro* spontaneous α-synuclein fibrils grown with concentrated monomeric α-synuclein in 50 mM Tris, pH 7.4, and 150 mM KCl over 168 hours of incubation described two novel classes of fibrils based on either inter-filament salt-bridges formed through K45 and E57 interactions (Polymorph 2a, shortened to “Class A” in this study), or a salt-bridge centered by amino acid E46 (Polymorph 2b, shortened to “Class B” in this study). The twisted DLB-CSF amplified fibrils from all ten samples segregate into these two filament bridge classes (**Table 2**). Samples amplified from DLB cases I, III, IV, VII and X are mixtures of Class A and Class B fibrils, whereas DLB cases VIII and IX have only Class A fibrils, and DLB cases II, V, and VI have only Class B fibrils (**Fig. 2**). These results underscore the physiological relevance of the Guerrero-Ferreira fibril classes in Lewy body dementia CSF^29^.

**Figure 2.**
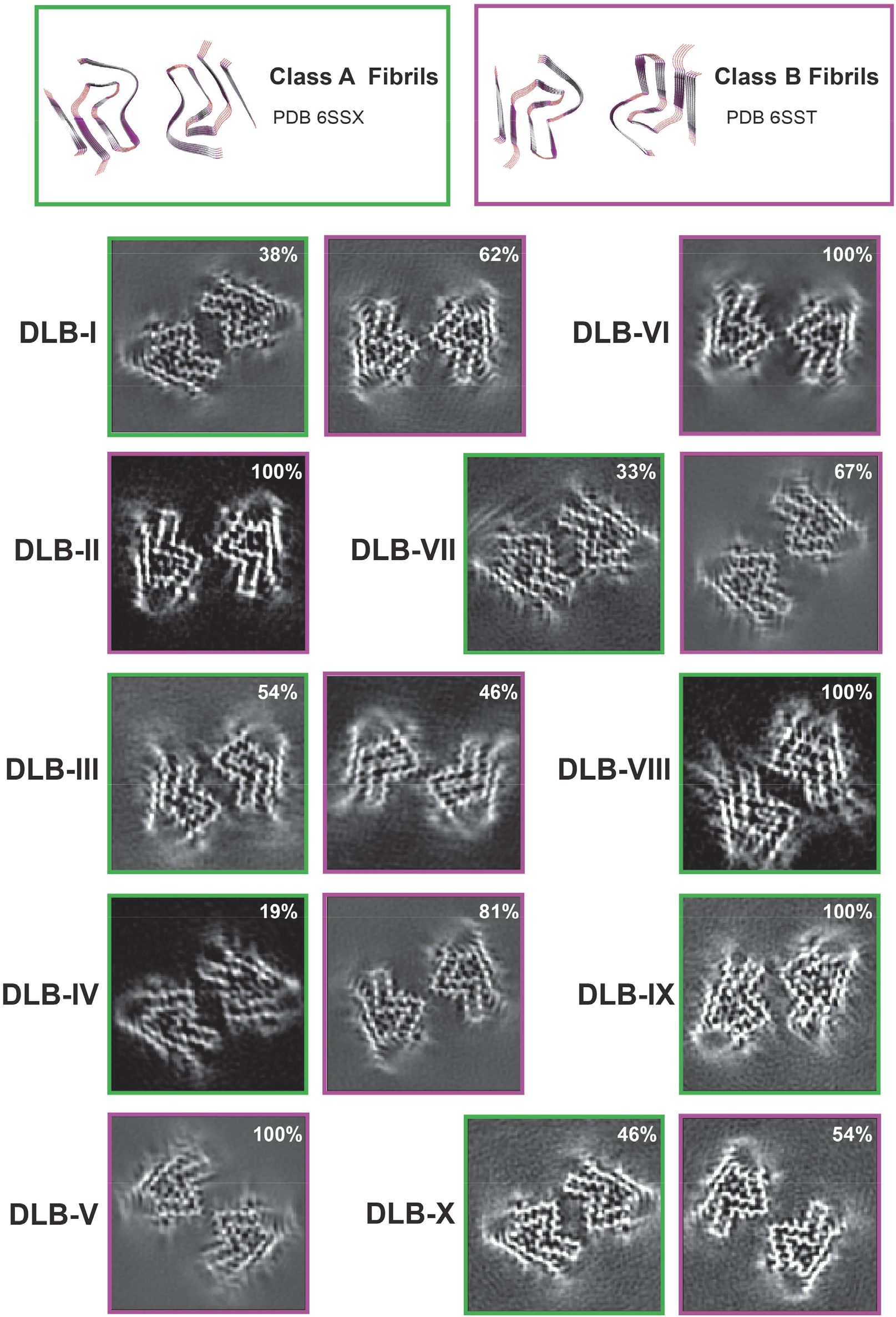
Projected fibril slices and their relative proportions from ten DLB-CSF amplified fibril products. Thicknesses (6 Å) from 3D cryo-EM maps are divided into two groups bearing similarity to two previously identified classes purified from *in vitro* spontaneous α-synuclein aggregation reactions under different salt and buffer conditions than those used here, named polymorph 2a (shortened to Class A here, 6SSX) and polymorph 2b (shortened to Class B here, PDB 6SST). 3D maps correspond to Fourier shell correlation plots (see **Supplemental Fig 2.**) and final resolutions (indicated in **Table 2**).

### High resolution cryo-EM structures of six novel α-synuclein fibril assemblies

All the protofilaments from the fibril products (DLB I-X) share similar folding features with β-strands from V15 to K96 in the WT full-length α-synuclein (**Fig 3a**). The eight β-strands form antiparallel pairs (i.e., β-strands 1:8, 2:7, 3:6, 4:5) that share a close interface with hydrophobic and hydrophilic interactions as graphically simplified in **Fig 3b**. Further 3D classification and helical reconstruction of the filaments associated with both Class A and Class B fibrils reveal differences between the fibril assemblies from different samples in the inner hydrophobic core arrangements among β-strands 1, 2, 7 and 8. The differences segregate into two possible hydrophobic core pocket models and are referred to here as “extended” or “compact”. The two pocket models share conserved β-strands 4-5 in a core stabilized by hydrophobic triple valine stacking: V52, V66 and V70, and hydrogen bonds: T54, T64, Q62, T59, and the main chain of A56. Similar fibrils core arrangements have been described (PDB 7NCJ, 7NCI, 7NCH, 7NCG, 7NCA, 6SSX, and 6UFR). However, compared to the conventional compact pocket arrangement, the extended arrangements found here flips β-strand 8 to glide along β-strand 5, which reorders the interactions by hiding F94 into the hydrophobic core thereby establishing a new salt bridge between E61 and K96, sealing the core. Further, β-strand 1, which is locked with β-strand 8 by the salt bridge between E83 and K21, follows along the gliding change. As a result, the disordered flexible loop (the cryo-EM density in this region is weak) between β-strand 1 and 2 shifts outwards, followed by β-strand 2 swinging into the same direction. Thus, β-strand 7 drags with β-strand 2 forming an enlarged hydrophobic pocket instead of shrinking with β-strand 6 (**Fig. 3c** and **Supplemental Fig.3**).

**Figure 3.**
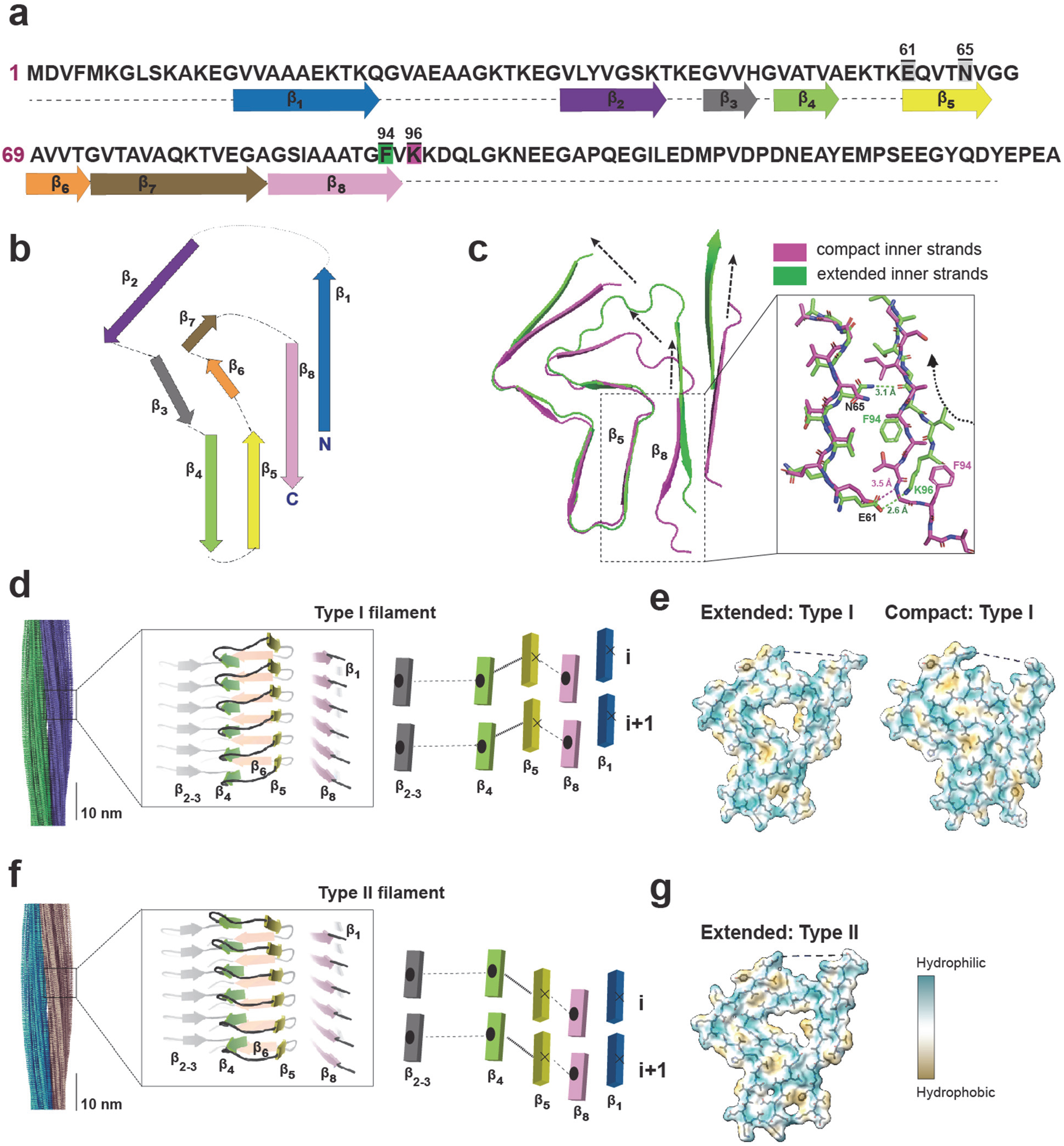
Identification of novel Type I and Type II β-sheet stacks in DLB-CSF amplified filaments. **a.** Primary sequence of human α-synuclein protein highlighting eight β-strands with colored boxes. In the primary sequence, key amino acids involved in Type I and Type II inner strand arrangement are color-shaded and highlighted with black bars. **b.** Generalized filament arrangement of β-strands 1 through 8 in the fibril products. **c.** Fitted atomic model of compact (observed only in Type I β-sheet stacked filaments) and extended (observed in both Type I and Type II β-sheet stacked filaments) that highlights (magnified black box) differential bonding caused by the compact or extended arrangements. **d,e.** 3D Cryo-EM fibril reconstructions with Type I filaments that position β5 above β4 found with both extended hydrophobic pockets and compact hydrophobic pockets. **f,g.** Type II filaments position β5 below β4 and always have extended hydrophobic pockets.

Further 3D classification reveals two markedly different configurations of β-sheet stacking as a result of the β-strand pairs spreading into different rungs. In left-handed helices formed by the triangular progressive rungs, β-strand 5, together with 6 and 7, rise up along the fibril axis, exemplifying a “Type I”β-stack arrangement (**Fig. 3d**). Both extended and compact hydrophobic cores are found with Type I β-stacking (**Fig. 3e**). However, in “Type II”β-stack arrangements, β-strand 5 and the following β-strand 6 and 7 drop below the fibril axis (**Fig. 3f**). Only extended pockets can be found with Type II β-stacking (**Fig. 3g**). All characterized DLB-CSF amplified fibril assemblies are thus described by fibril class A or B, extended or compact inner pocket configuration, and Type I or Type II β-stack arrangement. In this classification schema, five of the ten samples (DLB-II, V, VI, VIIII, IX) harbor a single fibril type. Two types of fibrils are identified in four of the ten samples (DLB-I, III, IV, X), and DLB-VII shows three types of fibrils. Final resolutions range from 3.8 Å to 2.9 Å, with an average twist pitch of 116 nm and a common stack interval of 4.8 Å (**Table 2** and **Supplemental Fig. 2,4**). The most observed fibril assembly, present in six of ten fibril product samples (PDB 8CYT in DLB-I and IV, PDB 8CZ2 in VII, VIII, IX, and PDB 8CZ6 in DLB-X), is a Class A, type I, extended pocket fibril (**Fig. 4**). Next most common, found in four of ten fibril products (DLB-I, IV, VI, and VII), is a class B, Type II, extended pocket fibril. A compact pocket fibril was found in only a single Type I assembly (DLB-III) with both class A and class B arrangements. This fibril type bears some resemblance to a previously described E46K-α-synuclein mutated fibril type^30^, with the E46K mutation causative for DLB^31^ but with WT-α-synuclein in this case (**Supplemental Fig. 5.**). Finally, a heterofilamentous mixed-type fibril assembly exists in both DLB-II and DLB-V and is composed of Type I and Type II filaments wound together with Class B salt-bridges.

**Figure 4.**
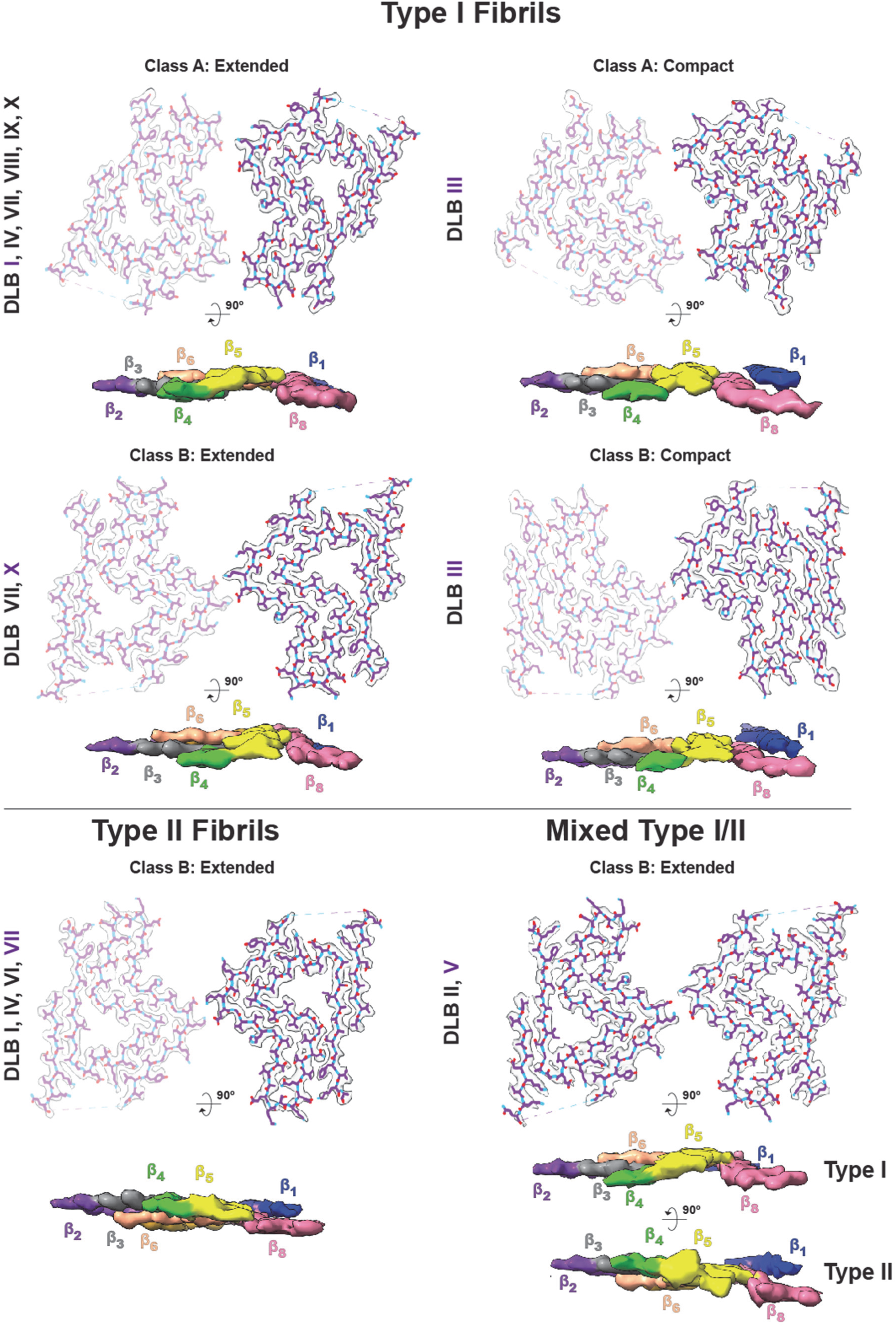
Cryo-EM reconstructions of six novel DLB-CSF amplified α-synuclein fibril strains. Novel fibril strains are differentially characterized based on Type I or Type II β-strand stacking, extended or compact pockets, and Class A or Class B filament-filament interaction. DLB-CSF samples that share the fibril strain assignment (β-strand type, fibril class, and extended/compact pocket) are indicated, with a purple-shaded font highlighting the DLB-CSF sample graphically depicted. Superimposed structural stick models in fitted cryo-EM densities show the amino group in blue and the acid group in red, with side-view arrangements immediately below depicting surface densities of the labeled β-strands.

### Amyloid dye binding characteristics of DLB-CSF amplified fibrils

As different types of hydrophobic amyloid dyes are known to bind to hydrophobic pockets and β-sheet stack assemblies specific to fibril conformations and no other conformations (e.g., monomeric), the different pockets and stacks revealed by cryo-EM might manifest with different amyloid dye binding profiles. Further, if the different structures are reliably amplified from the DLB-CSF samples, and not the product of stochastic nucleation and random structural propagation, then consecutive amplifications performed on different days from the same CSF sample should produce the same amyloid dye fingerprint in the resultant assemblies. To test these possibilities, we first created a large batch of reference bovine-serum albumin (BSA) fibrils to compare DLB-CSF fibril assemblies from amplifications from the original batch used for cryo-EM analysis as well as additional amplifications performed on a smaller scale. The BSA reference fibrils bound the hydrophobic amyloid dyes ThT, NIAD-4, and nile red (**Fig. 5a-c**). Calculated as a fold-binding to reference BSA fibrils, a diversity of DLB-CSF amplified fibril dye fingerprints emerge between the ten fibril products (**Fig. 5d-f**). While the DLB-I fibril product had similar ThT binding to BSA fibrils, the DLB-VII product had ~10-fold more ThT binding, whereas nile red and NIAD-4 binding was similar between DLB-I and DLB-VII samples. Variability in amyloid dye profiles was low between runs from different days, suggesting some level of conservation and non-stochastic structural features, although corresponding cyro-EM structures were not obtained with every amplified fibril assembly preparation. Raw fluorescent values for five independent amplifications performed in different weeks for DLB-VII amplified fibrils, with DLB-VII represented by a mix of three different types of fibrils, show minimal variability from amplification run-to-run (**Supplemental Fig. 6**). These results may be in line with previous reports that suggest α-synuclein fibril seeds from Lewy body disease stably propagate nuanced structural features in progeny fibrils.

**Figure 5.**
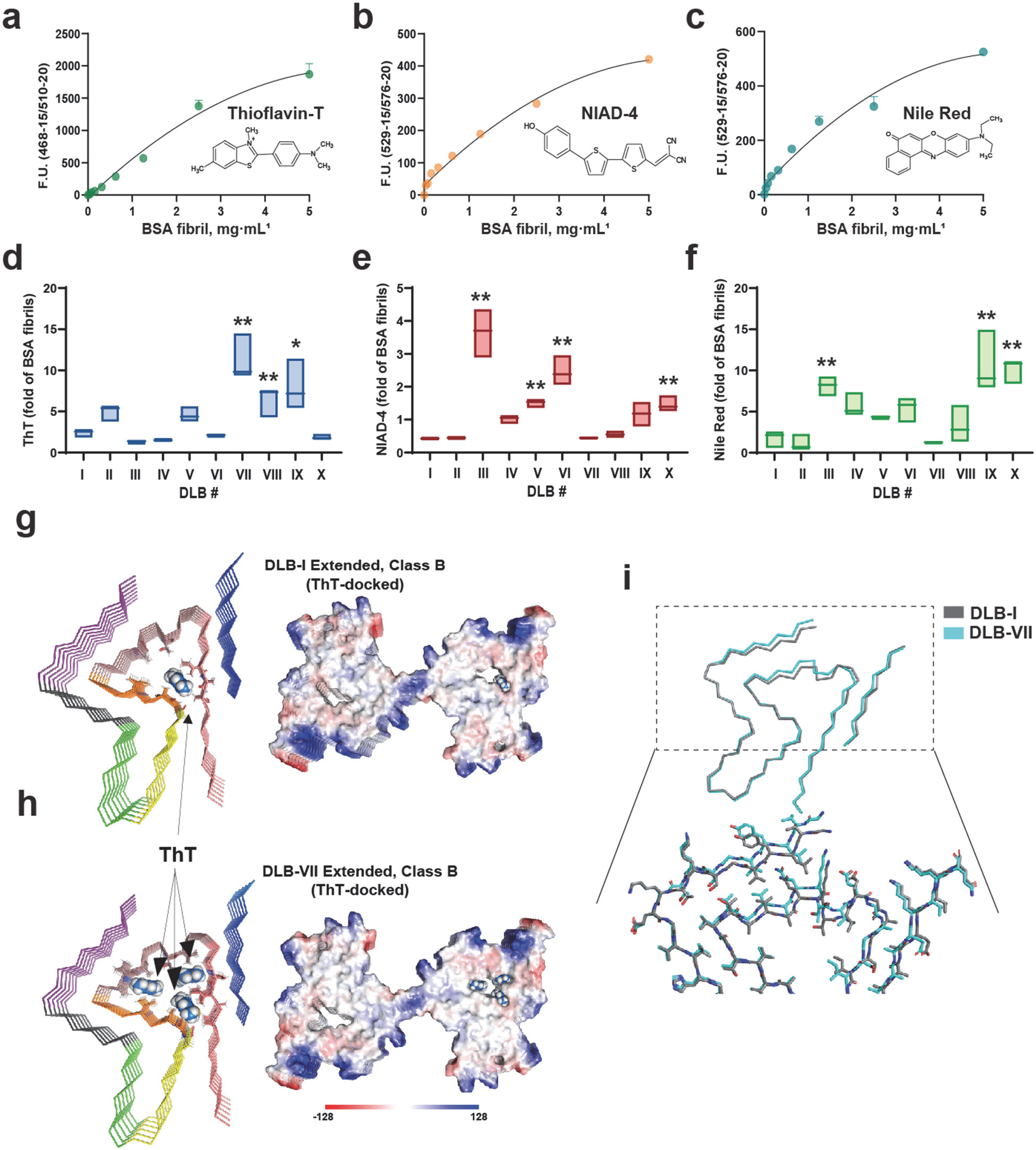
Distinct amyloid dye fingerprints of α-synuclein fibrils amplified from DLB-CSF. **a.** Representative reference binding curves for **a.** thioflavin-T (ThT) binding, **b.** NIAD-4 binding, and **c.** nile red binding to purified bovine serum albumin (BSA) protein fibrils. **d-f** Floating bars show the apparent mean, min, and max dye binding of three different amplifications of α-synuclein fibrils from the indicated DLB-CSF samples. * is p <0.05 and **is p<.005, oneway ANOVA with Dunnett’s post-hoc test (reference sample DLB-I). **g.** Predicted molecular docking of a ThT molecule in DLB-I Class B fibrils compared to **h.** Predicted ThT dye docking in DLB-VII Class B fibrils, with lines and electrostatics surface models shown. **i.** A magnified overlay of these two fibrils (DLB-I and DLB-VII, both Class B extended) highlight β-strand 6 and 7 offsets that alter the known ThT-high affinity hydrophobic binding pocket.

As the ThT amyloid dye is used extensively to characterize α-synuclein fibrils and in diagnostic aggregation assays for Lewy body diseases, we examined the cryo-EM structures associated with the highest ThT binding fibril preparation, DLB-VII, in comparison to one of the lower ThT binding fibril preparations, DLB-I. Specifically examining class B extended pocket assemblies, we defined a small 3D space around a hollow the hydrophobic region of both structures for glide docking analysis with Schrödinger software. DLB-VII fibril assemblies form three possible high affinity binding pockets to ThT in the NAC-region (β-sheets 5 to 8) while DLB-I exposes only a single ThT pocket in β-strands 5 to 8 (**Fig. 5g-i**). These results may explain why DLB-VII fibrils have high ThT binding and DLB-I fibrils have lower ThT binding. Consistently, DLB-III fibrils with the compact-hydrophobic pocket and poor ThT docking have poor observed ThT binding. However, DLB-III fibrils bind some of the highest levels of NIAD-4, ~10-fold more than the DLB-I fibrils. NIAD-4 is a dye with proposed specificity to amyloid-β fibrils^32^ and may involve interactions with flexible-domains. The hydrophobic-sensor dye nile red identifies the Type 1 extended-pocket fibrils in the DLB-IX and DLB-X samples with the highest binding, ~10-fold higher than DLB-I, II, and DLB-VII fibril assemblies.

### Dynamic and differential pro-inflammatory responses to novel DLB-CSF amplified fibril strains

Through largely unclear processes, α-synuclein fibrils or other seed-competent conformations can exit cells in disease, spreading into CSF or potentially interconnected cells in a prion-like manner. Protein fibrils associated with neurodegenerative diseases, including α-synuclein fibrils, are known to interact with microglia through different toll-like receptors, and increase MHC class II surface expression (e.g., HLA-DRα)^33–35^. Microglia HLA-DRα correlates with α-synuclein inclusions in Lewy body dementia^36^. To gain insight into whether the fibril products produced from DLB-CSF amplifications might differ with respect to evoked microglia phenotypes, we differentiated primary human monocytes procured from healthy volunteers to a microglia-like state as previously described^37,38^. Adherent cells here robustly express the microglia-specific receptor P2RY12 receptor, and within minutes the cells internalize exogenous DLB-CSF amplified α-synuclein fibrils (**Fig. 6a**). As different fibril sizes or lengths may influence cellular uptake in cells (i.e., in microglia or neurons), we carefully normalized fibril lengths between the different preparations with temperature-controlled sonication, monitoring product sizes with dynamic light scattering in achieving similar size distributions among the different preparations (**Supplemental Fig. 7**). We further ensured that the conjugation of the pH-sensitive pHrodo-dye to the different assemblies was similar between different fibril preparations (**Supplemental Fig. 7**). There was no apparent difference in the microglia uptake of the fibrils (**Supplemental Fig. 8**).

**Figure 6.**
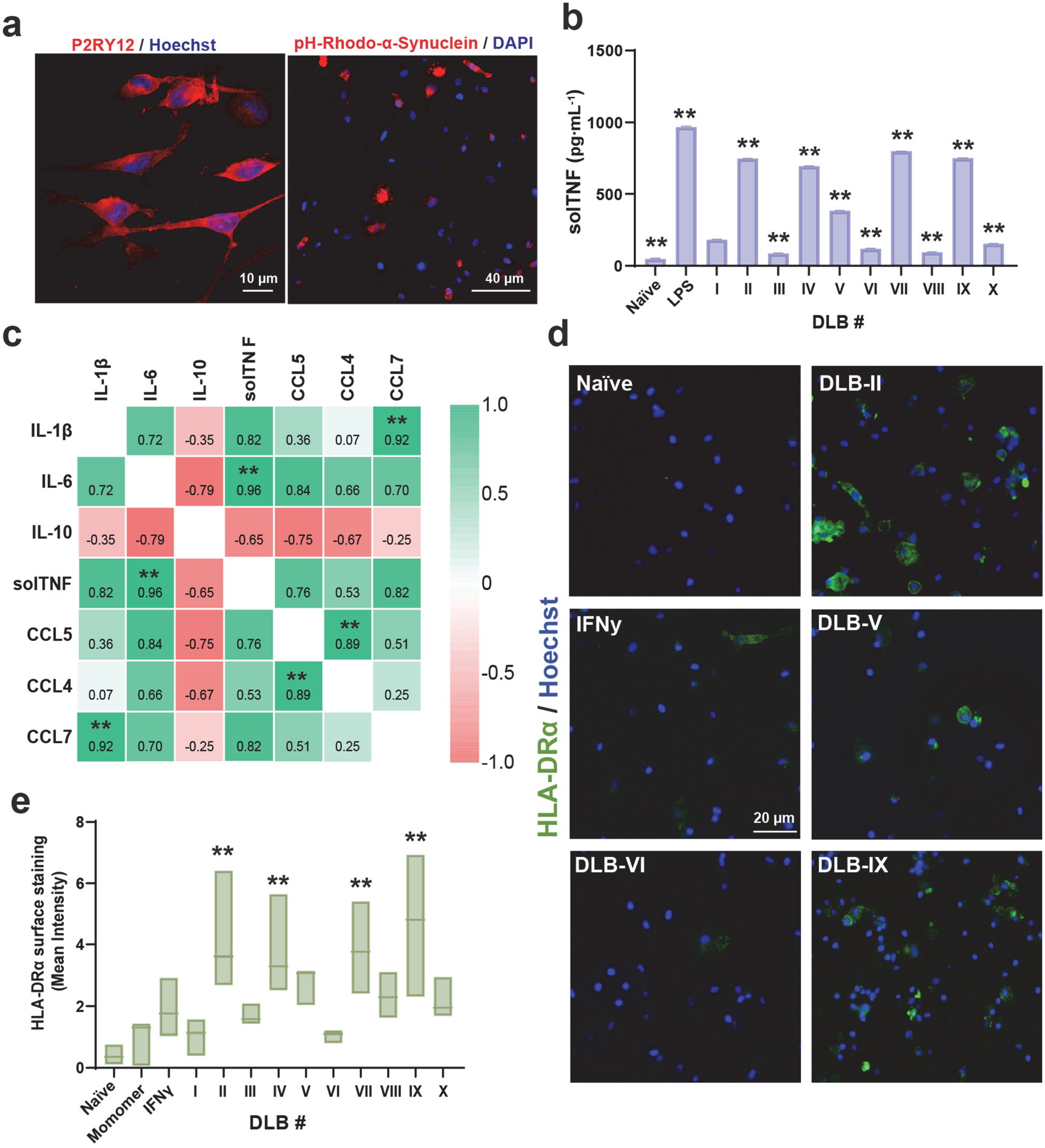
Different DLB-CSF amplified α-synuclein fibril mixtures cause profoundly different magnitudes of inflammatory responses in primary human microglia-like cells. **a.** Representative morphological characterization and strong reactivity of the human microglial-marker P2RY12, and strong cellular uptake of phRhodo-labeled α-synuclein fibrils two-hours after addition (1 μg per mL). **b.** Group quantification (ELISA) of soluble tumor necrosis factor (TNF) secretion six hours after fibril addition. Lipopolysaccharide (100 ng per mL) is included as a canonical positive control for TNF stimulation. **c.** Heat-map correlation of different measured pro- and anti-inflammatory secreted factors (ELISA) six hours after fibril stimulation. ** indicates significant correlation coefficients after correction for false discovery (p<0.0015, two-stage linear step-up, Benjamini, Krieger and Yekutieli). **.d.** Representative widefield immunofluorescence images of HLA-DRα surface expression (green) and Hoechst (blue) in microglia cultures intact or 24 hrs after fibril treatment, or interferon-γ (IFNγ) treatment (20 ng per mL) as a positive control for MHC-II induction.**e.** Floating bars show the apparent mean, min, and max HLA-DRα surface intensity from three different amplifications of α-synuclein fibrils from the indicated DLB-CSF samples.** is p<.005, one-way ANOVA with Dunnett’s post-hoc test (reference sample DLB-I).

Application of the same batch of amplified fibrils (i.e., DLB I-X) used for cryo-EM structural analysis to the microglia yielded marked differences in evoked TNF responses, exemplified by a ~10-fold increase in soluble TNF release between DLB-I and DLB-VII fibrils (**Fig. 6b**). This effect cannot be explained by potentially contaminating endotoxin associated with the recombinant fibrils, since the application of 1 μg per mL of fibrils applies negligible endotoxin (less than 0.001 E.U. per mL) exposures. Although physiologically relevant fibril concentrations are not clear, 1 μg per mL of amplified fibrils is a typical minimal seeding concentration of human fibrils used to induce aggregates *in vitro* in primary neurons days and weeks later^9,39,40^. Four of ten preparations of DLB-CSF amplified α-synuclein fibrils, DLB-II, IV, VII, and IX caused TNF secretion levels that approached 1 ng per mL soluble TNF, nearly matching the levels caused by high LPS exposures (**Fig. 6b**). In contrast, DLB-III, DLB-VI, DLB-VIII, and DLB-X caused minimal secretion of TNF that varied only 2-3-fold above basal TNF levels (i.e., naive, unstimulated). The DLB-III fibril sample, enriched with compact core Type I fibrils, elicited nearly ten-fold less soluble TNF production than extended core Type I fibrils enriched in DLB-IX fibrils. However, the DLB-VIII sample that is also enriched in extended core fibrils induced a minimal TNF section. The mixed Type I/II fibrils found in DLB II and V samples both caused elevated TNF secretion. In the evaluation of other cytokines and chemokines, pro-inflammatory soluble TNF levels were highly correlated with pro-inflammatory IL-1β (r=0.82, p<0.001) and IL-6 (r=0.96, p<0.001, **Fig 6c**). Unlike LPS, none of the preparations caused secretion of the proinflammatory IFNγ cytokine. The secretion of chemokines important in pro-inflammatory monocyte recruitment to tissues, CCL4 and CCL5, were likewise positively correlated with one another (r=0.89, p<0.001). One day after the addition of fibrils to the microglia, MHC-II (recorded with HLA-DRα surface expression, **Fig 6d,e**) significantly increased but was variable between fibril preparations and did not correlate with TNF secretion measured from the same cultures (r=0.38, p=0.27). Several of the fibril preparations, including DLB-I, III, and VI, and α-synuclein monomer protein exposures, did not cause a significant induction of MHC-II surface expression over basal (unstimulated) levels (**Fig. 6h,i**). In contrast, DLB-II, IV, VII, and IX caused a robust increase that went above and beyond the response caused by exposures to the canonical MHC-II-inducing cytokine IFNγ (20 ng per mL human IFNγ). In consideration of the cryo-EM structure analysis that was performed on the same batch of fibrils used to treat the microglia, we did not identify a particular structural feature or amyloid dye profile that accurately predicted pro-inflammatory responses. These results highlight striking variability in pro-inflammatory responses elicited by nominally different α-synuclein fibril assemblies.

### DLB-CSF amplified fibril strains drive differential seeding activity for new α-synuclein inclusions in neurons

Besides interactions with glia, extracellular and pre-formed α-synuclein fibrils are known to rapidly internalize into neurons to seed the formation of new phosphorylated (e.g., pS129) inclusions from endogenously expressed α-synuclein protein^9^. To avoid host-sequence interactions with mouse α-synuclein amino acid sequence that is different from human sequence in several critical β-strands segments^9,41^, we cultured primary hippocampal neurons from postnatal mice previously described^42^ that express human α-synuclein in lieu of endogenous mouse α-synuclein. Combining DLB-CSF amplified fibril strains (1 μg per mL) with mature neurons for seven days in culture results in the formation of cytoplasmic pS129-α-synuclein inclusions that do not occur with monomer treatment (**Fig. 7a,b**). Internalization of the different fibril preparations into the neurons, as estimated with pHrodo-dye conjugated fibril uptake six hours after exposure, is similar between all of the DLB-CSF amplified assemblies (**Supplemental Fig. 9**).

**Figure 7.**
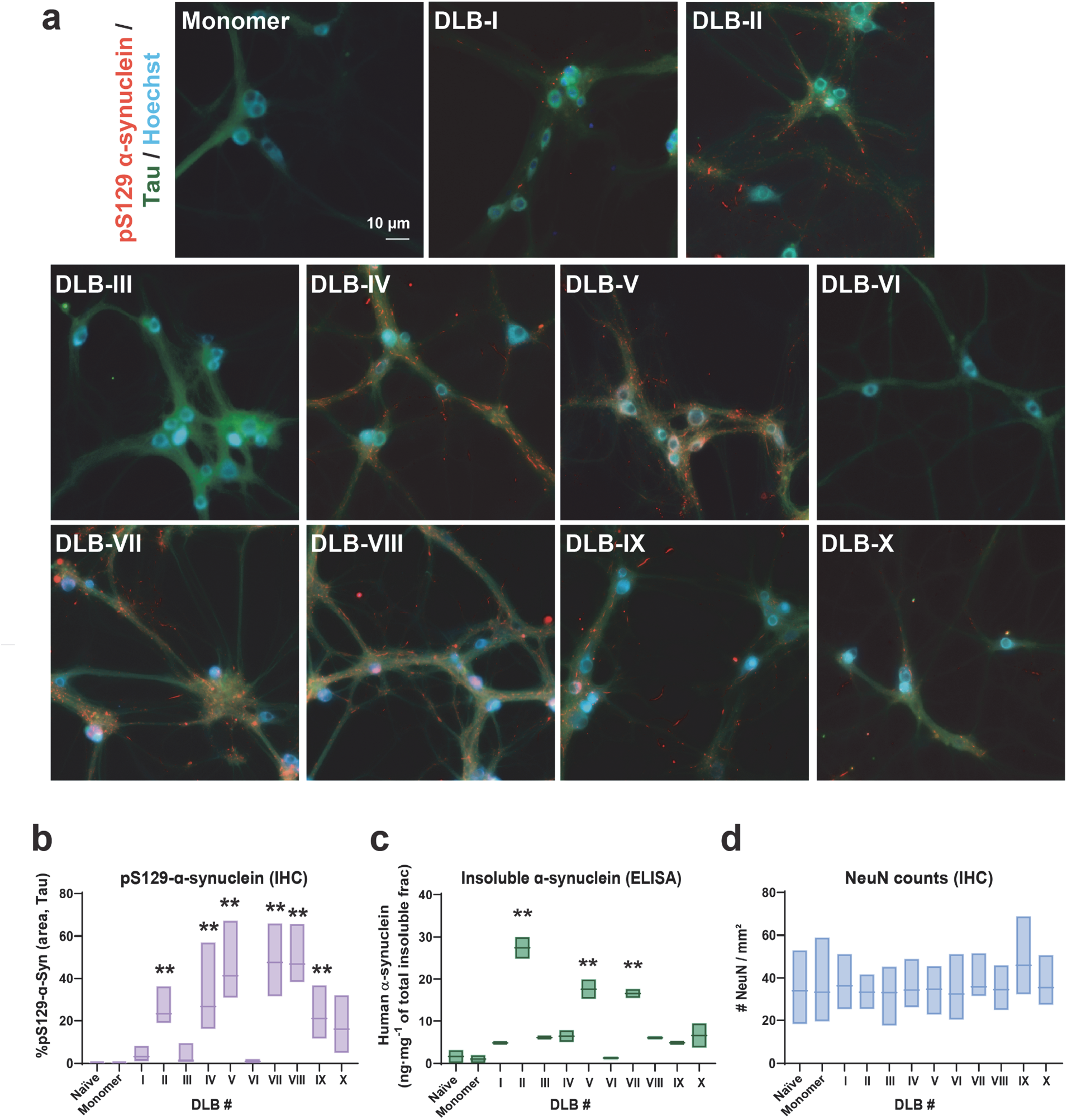
Differential neuronal seeding activity with DLB-CSF amplified α-synuclein fibrils. **a.** Primary neurons from SNCA^PAC-WT^ mice were cultured for seven days *in vitro* and treated with fibrils or monomer as indicated (1 μg per mL) for ten days before fixing and staining (immunofluorescence, pS129-α-synuclein) to determine the abundance of phosphorylated inclusions in tau-positive neuronal processes. Representative images show pS129-α-synuclein (red), tau (green) and Hoechst (blue). **b.** The percentage of tau area occupied by pS129-α-synuclein inclusions calculated from at least 20 images each from three independent cultures treated from three different fibril preparations of α-synuclein fibrils from the indicated DLB-CSF samples. **c.** The concentration of total α-synuclein deposited into the insoluble fraction of these cultures as measured by ELISA with monoclonal antibody 4B12, recognizing the 103-108 epitope. **d.** Corresponding NeuN-positive cell body counts as a function of survellied area in the cultures, each from three independent cultures treated from three different fibril preparations of α-synuclein fibrils from the indicated DLB-CSF samples. In these graphs, floating bars show the apparent mean, min, and max distributions from the indicated DLB-CSF samples, where ** is p<.005, one-way ANOVA with Dunnett’s post-hoc test according to the DLB-I reference sample. One-way ANOVA was not significant in panel d.

Applying the same batch of amplified fibrils used for cryo-EM structural analysis, pS129-α-synuclein inclusions appear ten days after fibril exposure but varied considerably in amount between the different fibril preparations (**Fig 7a**). DLB-I amplified fibrils, composed of class A and B, Type I and II fibrils, as well as the DLB-III fibrils, composed of compact-core Type I fibrils, had very low potency in the formation of new pS129-α-synuclein positive inclusions. In contrast, DLB-II, IV, V, and VII-X all had higher activity in seeding pS129-α-synuclein positive inclusions. To estimate the deposition of insoluble α-synuclein in a manner agonistic to phosphorylation at S129, in parallel cultures, neurons treated with fibrils for ten days were lysed first with a mild-detergent buffer, and then the pellet fraction sequentially lysed in SDS-buffer to access relatively insoluble α-synuclein. Diluting the SDS-insoluble fraction with saline and then measuring total α-synuclein with ELISA analysis, fibril preparations amplified from DLB-II, V, and VII again caused high depositions of α-synuclein into the SDS-solubilized protein fraction (**Fig. 7c**). Despite the high concentrations of insoluble α-synuclein formed in the cultures after fibril treatment (e.g., up to ~30 ng of insoluble α-synuclein per mg of total protein with DLB-II fibril treatment), there was no apparent loss of NeuN-positive cells (e.g., neurons) in the cultures in any of the fibril treatments at this time point (**Fig. 7d**). We did not identify any structural feature that would predict the observed seeding capacity differences in neurons and seeding responses in neurons did not correlate well with microglia pro-inflammatory responses from the same fibril preparations (**Supplemental Fig. 10**). Overall, these results highlight the striking landscape of structural and functional diversity associated with CSF-amplified α-synuclein fibrils from Lewy body dementia, and the apparent complexity of nuanced structural features that appear to cause large responses in neurons and microglia.

## Discussion

The novel results in this study center on four observations. First, fibrils amplified in the presence of DLB-CSF can be grouped broadly based on two distinct inter-filament classes apparent from 3D cryo-EM maps: Class A and Class B. Second, α-synuclein fibrils can be further classified based on two types of β-sheet rung stacking and two types of inter-β-strand arrangements that shape the hydrophobic pocket. The DLB-CSF amplified fibril products fall into six types of assemblies. According to amyloid dye-binding characteristics, which vary considerably between the different α-synuclein fibril assemblies, different rounds of amplification performed on different days and weeks produce fibril products with similar characteristics. Although cryo-EM structure analysis was not performed on multiple amplified fibril products performed on different days from the same CSF samples, extrapolating the observed low variability of the amyloid dye fingerprints in different rounds of amplification to the observed structural features that should be very sensitive to dye interactions suggests the fibril seeds from Lewy body disease CSF might allow for the stable propagation of core structural features. Third, the ten amplified fibril preparations studied here segregated into those that could cause high cytokine responses in microglia, or those that elicited weak responses, and those that could cause high MHC-II responses, or those with weak MHC-II responses. The responses caused by DLB-CSF amplified products varied more than 10-fold between some samples. The pro-inflammatory responses associated with some of the fibril preparations (e.g., DLB-VII) approached those of the canonical strong pro-inflammatory stimuli used as controls, whereas monomer equivalent protein was completely benign. Finally, in our fourth novel observation, the different fibril preparations varied considerably, more than 10-fold between some samples, in the seeding capacity in forming new inclusions and insoluble α-synuclein in mature primary neurons that express human α-synuclein. In consideration of all the quantitative variables measured here, a principal component analysis radially distributed the ten different fibril preparations according to PC1 and PC2 describing 42% and 21% of variance, respectively, highlighting the unique functional properties and structural features intrinsic to each combination of fibrils within the CSF-amplified fibril pools (**Supplemental Fig. 11**). Neither dyes, microglia responses, or neuronal responses could predict one another. These results implicate a previously undescribed intra-class fibril heterogeneity with a wide functional potency diversity that may be critical for functional phenotypes associated with these α-synuclein assemblies. Future studies with more CSF samples might be powered to assign disease features like progression and severity with observed structural and functional features associated with the CSF-amplified fibril products.

There are several important study limitations. First, we did not evaluate whether the amplified recombinant products (i.e., those fibril products permissive to amplification) match the structures of the parental seeds in the CSF, nor did we prove the parental CSF seeds are exclusively composed of α-synuclein protein. We have previously theorized based on a quantitative realtime α-synuclein aggregation monitoring assay that the concentration of small fibril seeds in CSF must be extremely low, in the femtomolar range or less, thereby precluding standard analytical purification and imaging^23^. In addition, it is possible that the fibrils that could propagate *in vitro* (i.e., amplify) were biased towards those that could amplify under the selected buffer conditions, and thus may not match the structural proportions of the starting seed compositions in the CSF. Further, whether or not the amplified recombinant products match the structure of inclusions present in the brain tissue of the associated CSF samples was also not determined here. Though, it is unclear how fibril seeds across a single brain might structurally differ in Lewy body dementia, and what particular brain regions might contribute the most fibril-forming seeds into CSF in the course of disease. Also, the circulating extracellular fibril seeds in CSF may not match the structure of large fibril depositions in Lewy pathology extracted by detergents, since the extracellular fibril fraction may be fundamentally different from fibrils able to escape the cell cytosol. Additional paired-sample matching cryo-EM studies comparing fibril structures obtained directly from different brain tissues from those amplified from CSF may be informative in this regard.

One recent study examining fibril twists with TEM analysis of fibrils amplified from formalin-fixed tissue extracts identified considerable structural variation within a single brain^43^. Our results may be consistent in that more than one type of fibril assembly was identified in most CSF samples. The most common type of α-synuclein fibril assembly identified, a class A, Type I, extended pocket fibril, found in more than half the samples analyzed, can structurally contrast with the rarest type of α-synuclein fibril identified, a compact core class A and B fibril found in only one sample. Interestingly, this rarer fibril assembly resembles the core structure of an α-synuclein fibril previously attributed to the pathogenic and very rare E46K mutation that is known to cause DLB. The DLB-III fibrils have relatively weak seeding activity in neurons and pro-inflammatory stimulation in microglia. PCA analysis highlights DLB-III with striking binding to the Aβ dye NIAD-4. Despite the weak ThT binding with DLB-III, the fibril products still bind enough ThT dye to monitor aggregation in real time, which speaks to the differential concentration of fibrils in disease that may be a critical component in disease phenotypes. In our functional studies, we normalized fibril concentrations prior to the addition to cells, but in disease, fibril concentrations may be an important variable that affects disease outcomes. Physiological concentrations of fibrils may vary such that high concentrations of weak fibrils may phenocopy high potency but lowly-concentrated assemblies of α-synuclein in disease.

Notably, our described approach for CSF analysis of fibril assemblies may not be broadly applicable across all α-synucleinopathies. In MSA brain tissue, it is unclear how well amplified fibrils resemble those of the fibril seeds extracted from tissue^27^, with MSA fibrils showing considerable differences in structure and function compared to those fibrils from Lewy body diseases^44^. Indeed, MSA fibrils apparently are not permissive to amplification from CSF in several types of aggregation assays^26,27^, and so the approaches described here may not be feasible to explore the structural diversity of fibrils from CSF from MSA cases. However, it seems possible that *in vitro* conditions may be identified in the future that permits the amplification of MSA fibrils that better resemble authentic assemblies purified from MSA brain tissue.

Overall, our results suggest that no single parameter studied here was successful in predicting the functional potency or underlying structural compositions and heterogeneities associated with α-synuclein fibrils amplified from DLB-CSF. Rather, nuanced structural changes in heterogeneous blends of fibril assemblies may underlie large functional differences. Tailored anti-α-synuclein therapeutics aimed at mitigating the effects of these different pathogenic species may be needed for successful interventions in disease.

## Data Availability

The cryo-EM maps have been deposited to the Electron Microscopy Data Bank (EMDB) under the accession codes EMD-27082 (spont.), EMD-27083 (DLB-I class B type II), EMD-27084 (DLB-I class A type I), EMD-27085 (DLB-II class B mixed), EMD-27086 (DLB-III class B type I), EMD-27087 (DLB-III class A type I), EMD-27088 (DLB-V class B mixed), EMD-27089 (DLB-VII class B type I), EMD-27090 (DLB-VII class B type II), EMD-27091 (DLB-VII class A type I), EMD-27092 (DLB-X class B type I), EMD-27093 (DLB-X class A type I). The coordinates of the corresponding models have been deposited to the Protein Data Bank (PDB) under accession codes 8CYR (spont.), 8CYS (DLB-I class B type II), 8CYT (DLB-I class A type I), 8CYV (DLB-II class B mixed), 8CYW (DLB-III class B type I), 8CYX (DLB-III class A type I), 8CYY (DLB-V class B mixed), 8CZ0(DLB-VII class B type I), 8CZ1 (DLB-VII class B type II), 8CZ2 (DLB-VII class A type I), 8CZ3 (DLB-X class B type I), 8CZ6 (DLB-X class A type I). Other data are available from the corresponding author upon reasonable request.

## Acknowledgements

This research was funded by NIH/NINDS R01-NS064934 (A.B.W), P50-NS108675 (A.B.W), and in part by Aligning Science Across Parkinson’s [ASAP-020527] through the Michael J. Fox Foundation for Parkinson’s Research (MJFF). For the purpose of open access, the author has applied a CC BY-ND public copyright license to all Author Accepted Manuscripts arising from this submission. Cryo-EM image collection work was performed in part at the Duke University Shared Materials Instrumentation Facility (SMIF), a member of the North Carolina Research Triangle Nanotechnology Network (RTNN), which is supported by the National Science Foundation (award number ECCS-2025064) as part of the National Nanotechnology Coordinated Infrastructure (NNCI). This study utilized the computational resources offered by Duke Research Computing (http://rc.duke.edu). We thank C. Kneifel, K. Kilroy, M. Newton, V. Orlikowski, T. Milledge, and D. Lane from the Duke Office of Information Technology and Research Computing for providing assistance with the computing environment.

## Methods

### Mouse strains and human samples

All study protocols were approved by the Duke University Institutional Review Board and the Institutional Animal Care and Use Committee. Post-mortem cerebrospinal fluid samples obtained from the Duke University Bryan Brain Bank are described in **Table 1** (additional information for these samples including α-synuclein seeding activity have been described^23^). DLB-CSF samples are from pathologically confirmed neocortical-type dementia with Lewy bodies, or a healthy control, and were processed as coded samples, with investigators blinded to group identity until final data curation. Mice were housed under environmentally controlled standard conditions with a 12-hr light/dark cycle and free access to food and water. Strains used include WT-hPAC-α-synuclein mice (SNCA^PAC-WT^); Snca^-/-^ (Jax Inc, RRID:IMSR_JAX:010710), that express endogenous levels of human α-synuclein and no mouse α-synuclein, and knockout-α-synuclein mice (Snca^-/-^) equivalent on the same strain that lacks any α-synuclein expression^45^. Females and males were equally distributed and randomized in groups throughout the study.

### Generation of monomeric α-synuclein

*Escherichia coli* BL21 (DE3) CodonPlus cells (Clontech, Inc.) were transformed with human α-synuclein-encoding plasmid (pET-21a(+)) and plated on Luria broth (LB) agar containing ampicillin (50 μg per mL). Protein expression in bacterial cultures was induced at an optical density of 0.8 with 0.05 mM IPTG (RPI, Inc.). Collected pellets were lysed in 750 mM NaCl, 10 mM TrisHCl, pH 7.6, 1 mM EDTA, 1 mM PMSF, and sonicated at 70% power (Fisher500 Dismembrator, Fisher Inc.) for 1 min each 30 mL aliquots. Homogenates were boiled at 90°C for 15 minutes, cooled on ice, and centrifuged at 20k *xg* for 30 min. Supernatants were passed through a 0.45 μm vacuum filter (Thermo-fisher, Inc.) and dialyzed with 25 mM NaCl, 10 mM Tris, pH 7.6, and 1 mM PMSF. α-Synuclein protein was extracted via a HiPrep Q HP 16/10 column (Cytiva, Inc.) for anion-exchange chromatography on an Akta Pure protein purification system (Cytiva, Inc.). Collected α-synuclein fractions assessed through SDS-PAGE and Coomassie stain were dialyzed and concentrated with a 3K M.W.C.O. filter (Amicon, Inc.) and subjected to a second round of anion-exchange chromatography as previously described. Selected fractions assessed for purity by SDS-PAGE and Coomassie blue staining were subjected to second rounds of dialysis and concentration, and endotoxin levels were then reduced through multiple passes through endotoxin removal columns (GenScript, Inc.) to achieve contaminating endotoxin units below 0.1 per mg of protein, as measured through LAL chromogenic endotoxin quantification (GenScript, Inc.). Protein concentration was determined by UV absorption at 280 nm using a nanodrop system (Thermofisher, Inc.) and size distribution for monomeric forms was monitored via dynamic light scattering (DLS) on a Titan DynaPro (Wyatt Technology, Inc.) at 25°C with Dyna V6.3.4 software.

### Preparation of α-synuclein fibrils amplified from CSF

Solutions of 5 mg per mL monomeric α-synuclein in phosphate-buffered saline (PBS) were combined with 10% CSF (w/v) and incubated for up to 60 hrs with 1 min on and off cycling of 1,000 R.P.M. at 37°C. CSF samples were previously characterized for α-synuclein seeding activity via quantitative real-time quaking-induced conversion assays. Selected CSF samples for this study were from Lewy body dementia cases (i.e., neocortical-type DLB) with known high-seeding CSF, intermediate seeding, low seeding, and a control CSF sample with no detectable seeding activity ^23^. After 60 hrs, suspensions of white precipitated fibril layers from the reactions with CSF from DLB cases were centrifuged at 20k xg at room temperature to remove supernatants, and fibril pellets were washed three additional times in PBS. At 60 hrs of incubation, incubations with control CSF, or reactions without any CSF, did not yield detectable fibril products. To determine fibril concentrations, heavy fibril aliquots were titrated into 3 M guanidine chloride and incubated with agitation at room temperature for 5 minutes. Time-dependent DLS measurements were performed to ensure all fibril species returned fully to monomeric states, and then concentrations were determined through A280 measurements. Fibril assemblies were next analyzed via transmission electron microscopy at an applied concentration of 0.4 mg per mL, with fibrils absorbed to glow-discharged 300 mesh (Ladd Research Industries, Inc.) and stained with 2% uranyl acetate (Electron Microscopy Sciences, Inc.) in water. Images were acquired with an FEI Tecnai F20 electron microscope (Eindhoven, Inc.) operated at 80 kV with nominal magnifications at 30,000x and 66,000x and a defocus range of −1.0 μm to −1.27 μm. Typical fibril lengths of >1 μm were resolved in image analysis. For some experiments, fibril preparations were labeled with 80 μg of pHrodo iFL STP (P36010, Invitrogen) at 1 mg per mL concentration in 100 mM sodium bicarbonate overnight. Unconjugated dye was removed through consecutive rounds of pelleting heavy fibrils via low-speed centrifugation and washing in PBS. Prior to application on neurons or intracranial injections, fibrils were partially sonicated to matched lengths of 30-40 nm with polydispersity indices <0.5 as determined by dynamic light scattering assays the same day as experimentation. Fibril sonication was achieved with a cuphorn sonicator in thin-walled 0.2 mL tubes (Q-Sonica, Inc).

### Cryo-electron microscopy and helical reconstructions

Amplified fibril strains generated as described above were subjected to a brief water-bath sonication to disrupt larger aggregates for ~1 min in a water bath sonicator (Q-Sonica, Inc) and applied to plasma-cleaned (Pie Scientific) 1.2/1.3 UltrAufoil (Quantifoil) grids with a Leica GP2. Images were acquired on a Titan Krios Cryo-TEM at a dose of 60 electrons per A^2^ using Latitude data collection procedures (Gatan, Inc.). Image alignments and fine alignments were performed prior to the atlas collection. **Table 2** outlines selected data collection parameters. Movie frames were gain-corrected, aligned, and dose-weighed using Relion 4.0 software^46^. Contrast transfer function (CTF) parameters were estimated using CTFFIND-4.1 software^47^. For helical reconstructions, ~40 images were selected randomly for manual picking and ~14k fragments were extracted with inter-particle distance of 6 β-rungs of 28.5 Å (with estimated helical rise of 4.75 Å) from the start-end coordinate pairs. Extracted coordinates were used for training CRYOLO^48^. The same model was used for all image datasets in auto picking mode with CRYOLO. Fragments were first extracted using a box size of 1024 pixels and down scaled to 256 pixels resulting in a pixel size of 4.32 Å. Reference-free 2D classifications were performed to assess different fibril strains, cross-over distances (estimated as 1,200 Å), and helical rise (4.82 Å), and to select suitable fragments for further processing. Initial references were generated from selected 2D class averaged images using the Relion_helix_inimodel2d command using a cross-over distance of 1200 Å, search shift of 15 Å, and search angle of 15 degrees. The clean set of fragments was re-extracted using box sizes of 512 pixels and down-scaled to 256 pixels (binned pixel size of 2.16 Å) for optimal performance in 3D classifications. To minimize reference bias, initial models were low-pass filtered to 40 Å. 6-8 classes were used during 3D classification together with initial helical twists set to −0.7 degrees, initial helical rises of 4.82 Å, and regularization parameter T of 4. Helical symmetry local search was performed with helical twist ranges varying from −0.85 to −0.5 and helical rise ranges varying from 4.8 to 4.84 during classification. Only the central 10% of the z length was used for reconstructions. Classes with nominal folding features were selected individually and re-extracted with box sizes of 384 pixels. Additional rounds of 3D classification were performed with the new reconstructions as reference using an initial low-pass filter set to 10 Å to clean the particles further. Classes with the same fibril types and folding features were merged together for 3D auto refinement. Initial helical twist and helical rise parameters were updated from 3D classifications. Local symmetry search was not performed at the first round of refinement. In the second round refinement, a shape mask comprising 90% of the central z length was applied along with local symmetry search. To improve the resolution, multiple rounds of CTF refinement, Bayesian polishing, and refinements with previous results using a particle extraction size of 256 pixels were used. For strains with high P21 screw symmetry, an additional round of refinement with the screw symmetry applied was conducted. Post processing was accomplished using a shape mask comprising 10% of the central z length, and final resolutions were estimated for each dataset using the gold standard FSC 0.143 cutoff.

### Fibril modeling

Two dimerization related chains from PDB IDs 6SST and 6SSX were used as initial models. Residues fitting model geometries were refined manually using COOT based on each sharpened density map^49^. Helical symmetry parameters from reconstructions were applied to the templates to generate β-strand stacks with 6 or 7 rungs. Final models were refined in PHENIX using phenix.real_space_refine (Phenix-1.2-4459)^50^. Manually defined non-crystallographic symmetry constraints according to the helical symmetry and β-sheet secondary structure restraints were imposed during refinement. Multiple iterations of PHENIX refinement and manual adjustment with COOT were performed and model evaluation was done using Molprobity, **Table 2**^51^.

### Docking

Molecular docking studies were performed using the Schrödinger Glide docking method^52–54^. The ThT coordinates were extracted from PDB ID 3MYZ. The docking grid was restrained within the protofilament hydrophobic core pocket surrounded by β-strands 6, 7, 8. Considering the depth of the pocket was formed by repeated rungs, 6 rungs were included in the grid, leaving enough gliding space for the ThT pose searching. Hydrophobic interactions weights were elevated during scoring based on the fact that ThT and the pocket are strongly hydrophobic. 3 rounds of prediction were performed for each target to validate the consistency of the results.

### Amyloid-dye assays

Fresh aliquots of amyloid-binding dyes, including 15 mM NIAD-4 (Caymanchem, Inc.), Nile Red (Sigma, Inc.), and 1 mM Thioflavin-T (Sigma, Inc.), were prepared in DMSO and sequentially diluted into saline at a concentration of 200 μM and added to ultra-low binding 384-well plates with clear bottoms (Corning, Inc.). For terminal dye binding estimates, fibril products were combined to 0.2 mg per mL and fluorescence intensity recorded with excitation/emission spectra set at 529-15 nm/576-20 nm for NIAD-4 and Nile Red, 448-10/482-10 nm for Thioflavin-T on a Clariostar Omega reader (BMG, Inc). Standard curves of varying concentrations of high-molecular weight albumin (BSA, Sigma, Inc.) were generated for each run. For kinetic dye measurements during fibril growth, in amplification assays described above, samples were supplemented with 10 μM Thioflavin-T (Sigma, Inc.) in ultra-low binding 384-well plates (Corning, Inc.) with clear bottoms and sealed with foil (BioRad, Inc.). Each plate was supplemented with a standard curve of serial dilutions of spontaneous α-synuclein reference fibril particles as described^23^. Reaction fluorescence was monitored at 448-10 nm excitation and 482-10 nm emission on FLUOstar Omega (BMG, Inc.) plate reader every 30 minutes with intermittent shaking at 700 R.P.M.

### Primary mouse neuron cell culture and analysis

Post-natal (P0) SNCA^PAC-WT^;Snca^-/-^mice, with human α-synuclein in place of mouse α-synuclein as described^23,42^, were incubated on ice for five-minutes. Their brains were removed, and the hippocampi were dissected into Hibernate E medium (BrainBits, Inc.). Tissue was triturated into a papain solution (Worthington Biochemical, Inc.) in HBSS buffer (Sigma, Inc.) supplemented with 10 mM HEPES pH 7.4, 100 mM sodium pyruvate, and 1% penicillin/streptomycin (all from Thermo-Fisher, Inc.). Cells were plated into Neurobasal media supplemented with B-27, 5 mM GlutaMAX (all from Thermo-Fisher, Inc.), and 10% fetal bovine serum (Atlanta Biologicals, Inc.) onto poly-d-lysine (Sigma, Inc.) coated wells at a density of 1×10^5^ cells per cm^2^. 24 hours after plating, media was exchanged with Neurobasal media supplemented with GlutaMAX and B27 supplement. At day-in-vitro (DIV) 10, fibril strains were added to each well of neurons to a final estimated concentration of 0.62 nM (1 μg per mL). At endpoints, neuron plates were treated with 2% paraformaldehyde (Electron Microscopy, Inc.) in PBS for 15 min, washed, and incubated for 30 min at room temperature in 3% normal goat-serum (Equitech-Bio, Inc.) with 0.1% saponin (Sigma, Inc.) in PBS. Antibodies were then added to 1% goat-serum, 0.02% saponin buffer, in PBS. Primary antibodies include pS129-α-synuclein antibody (20 ng per mL, clone EP1536Y, RRID:AB_765074, Abcam, Inc.) and anti-Tau (0.5 μg per mL, clone Tau-5, RRID:AB_1087687, Invitrogen, Inc.) incubated overnight at 4°C. Neurons were rinsed, and then incubated in a solution of secondary antibodies, including goat anti-rabbit AlexaFluor 555 and goat anti-mouse AlexaFluor 647, and DAPI stain (1 μg per mL, Thermo-Fisher, Inc.) for 3 hours at room temperature. Images were obtained and analyzed by investigators blinded to sample identity on a Keyence BZ-X810 microscope using the manufacturer’s software.

### Primary human microglia and analysis

Blood mononuclear cells were isolated from venous blood draws from healthy volunteers and processed with SepMate (Stemcell Tech, Inc.) tubes and Lymphoprep (Stemcell Tech, Inc.) tubes as described^38^. EasySep Negative Selection Human Monocyte Enrichment kits (Stemcell Tech, Inc.), without CD16 depletion, were used according to manufacturer’s instructions. Adherent cells were cultured into DMEM (Invitrogen, Inc.) supplemented with Glutamax (Invitrogen, Inc.), 10% fetal bovine serum, 1% penicillin/streptomycin (Invitrogen, Inc.), Fungizone (2.5 ug per mL), M-CSF (10 ng per mL), GM-CSF (10 ng per mL), NGF-β (10 ng per mL), CCL2 (100 ng per mL), and IL-34 (100 ng per mL) as described^37^. All cytokines were purchased from PeproTech. Cells were cultured for 12 days before experiments to obtain microglia-like cells with transcriptional signatures resembling resident brain human microglia^23,37^. As indicated, some cells were activated with LPS (100 ng per mL, triple-purified from E. coli strain O55:B5, InvivoGen, Inc.) or IFNγ (20 ng per mL). Live primary human microglia (Day 12) were washed with endotoxin-free PBS (Sigma, Inc.) three times and put on ice for 10 mins. Then the live microglial cells were incubated with PBS supplemented with 5% BSA + HLA-DR mouse monoclonal antibodies pre-conjugated with FITC (1:100, BioLegend, Inc.) + Hoechst 33342 (1:5000, Thermo-Fisher, Inc.) for another 10 mins on ice. After washing with endotoxin-free PBS on ice three times, the live microglial cells were fixed with 4% PFA.

### Enzyme-linked immunosorbent assays (ELISA)

Soluble and secreted chemokines and cytokines were analyzed via DuoSet ELISA systems (R&D, Inc.) according to manufacturer’s instructions and include human IL1β, IL-6, IL-10, soluble TNFα, CCL5, CCL4, and CCL7. Quantification of human α-synuclein concentration in SDS-solubilized fractions of primary hippocampal cultures was performed according to manufacturer’s instructions with the LEGEND MAX™ platform with the monoclonal antibody 4B12, binding the epitope 103-108 still present in C-terminal truncated forms of α-synuclein (Biolegend, Inc.). In brief, treated cells were lysed in 0.1% Triton X-100 with phosphate and protease inhibitors (Roche, Inc.) and the resulting pellet after high-speed centrifugation was then dissolved in 1% Triton X-100, 0.1 % SDS in PBS. Pellets were further solubilized by probe-tip sonication at 10% power for 10 sec and 1:400 dilutions were applied onto ELISA plate along with serial dilutions of recombinant α-synuclein fibrils prepared under the same conditions. Chemiluminescent signal was recorded on a ClarioStar plate reader (BMG, Inc.) and an extracted equation from a standard curve generated from serial fibril dultions was applied to identify the level of α-synuclein in cell lysates.

### Statistical analysis

Statistical analyses were conducted using the GraphPad Prism 9 software. Specific statistical tests for each dataset are described in each figure legend.

## Supplemental Figures

**Supplemental Figure 1.**
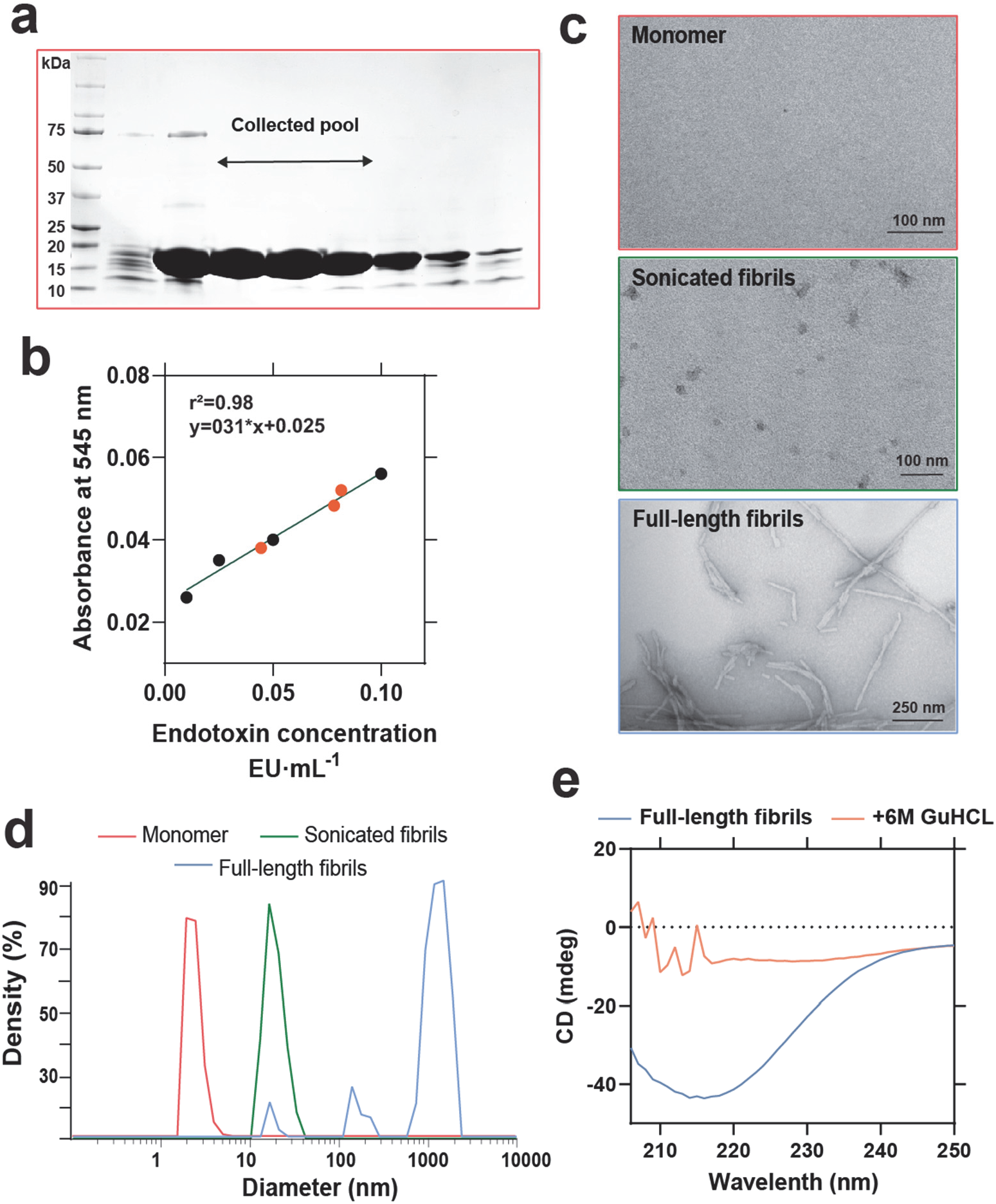
Validation and quality analysis of α-synuclein monomeric protein used for fibril amplifications. **a.** Representative Coomassie blue staining of an SDS-PAGE gel highlighting the collected pools of pure (estimated >98%)α-synuclein monomer fractions eluted with anion-exchange chromatography on HiPrep Q HP 16/10 columns. **b.** Representative standard curves of endotoxin quantification of α-synuclein preparations after endotoxin depletion. Black data-points show known endotoxin standards against high-concentration (70 μM in the assay) α-synuclein monomer protein, with results from the three batches of protein used in this study shown. Levels of remaining endotoxin were measured at 0.05-0.1 endotoxin units (EU) per milligram of protein using the LAL method. **c.** Representative negative electron scan micrographs of monomer, sonicated, and full-length fibrils, along with **d.** representative histograms of molecular diameter relative to percent of density acquired by dynamic light scattering measurements. **e**. Circular dichroism spectra of full-length fibrils and after 5 min incubation in 6M guanidine chloride (GuHCL). 6M GuHCL treatment was always used prior to estimation of protein concentrations via A_280_.

**Supplemental Figure 2.**
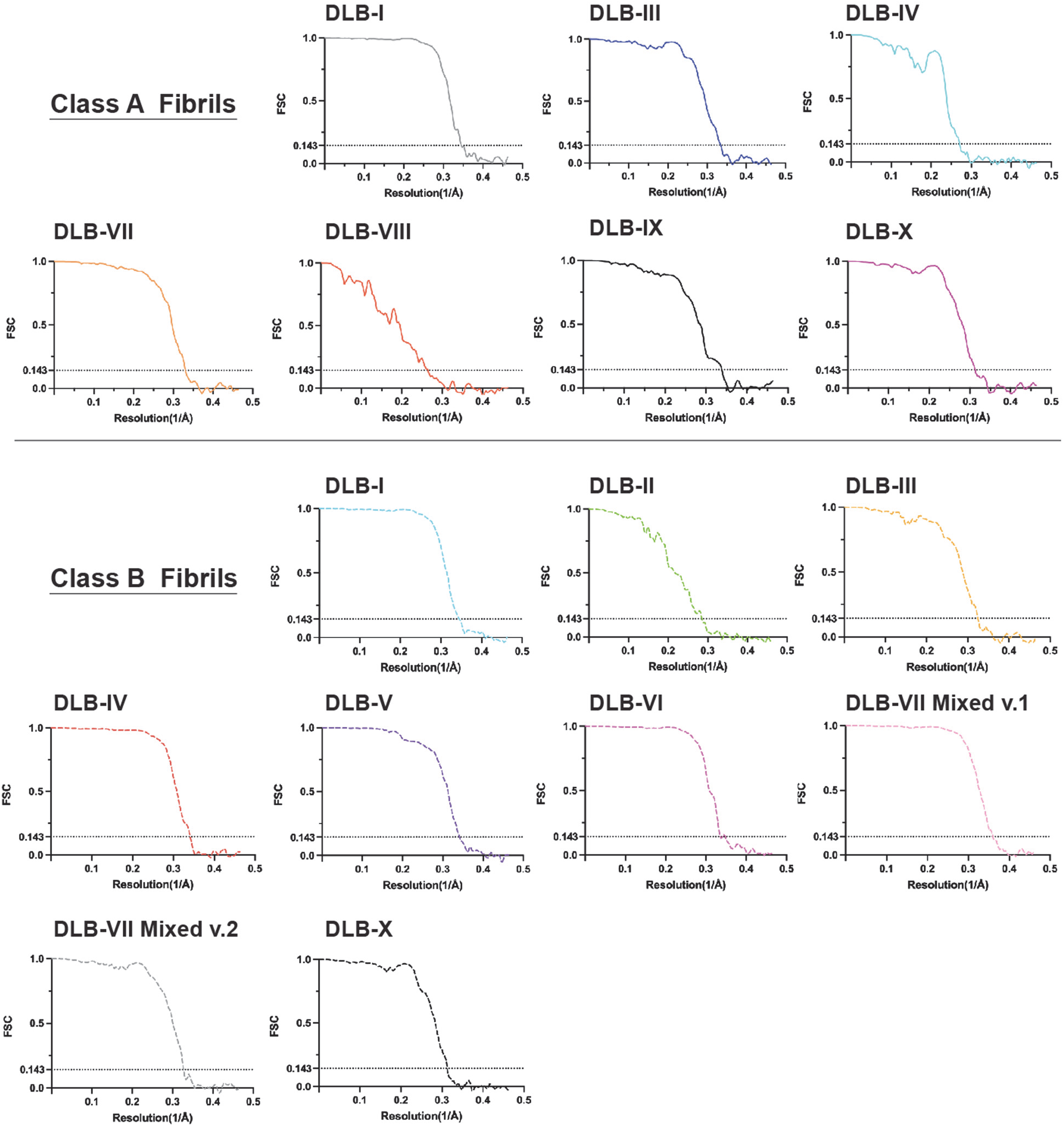
Resolution estimation and validation of Cryo-EM collected images of procured α-synuclein fibril strains. Fourier shell correlation (FSC) for the 3D reconstruction of the cryo-EM maps. Class A and B fibril strains were identified through 2D reconstruction and the majority of the DLB-derived fibril preparations contain two populations with distinct structural arrangement. DLB-III is referred to class A accordingly and a lower resolution estimation for both cases is associated with flexibility of the structure. DLB-VII Class B (v.1 and v.2) are associated with two different assemblies with the same 3D map (**Fig 2**). Detailed collection data for the cryo-EM analysis with final resolution is described in **Table 2**.

**Supplemental Figure 3.**
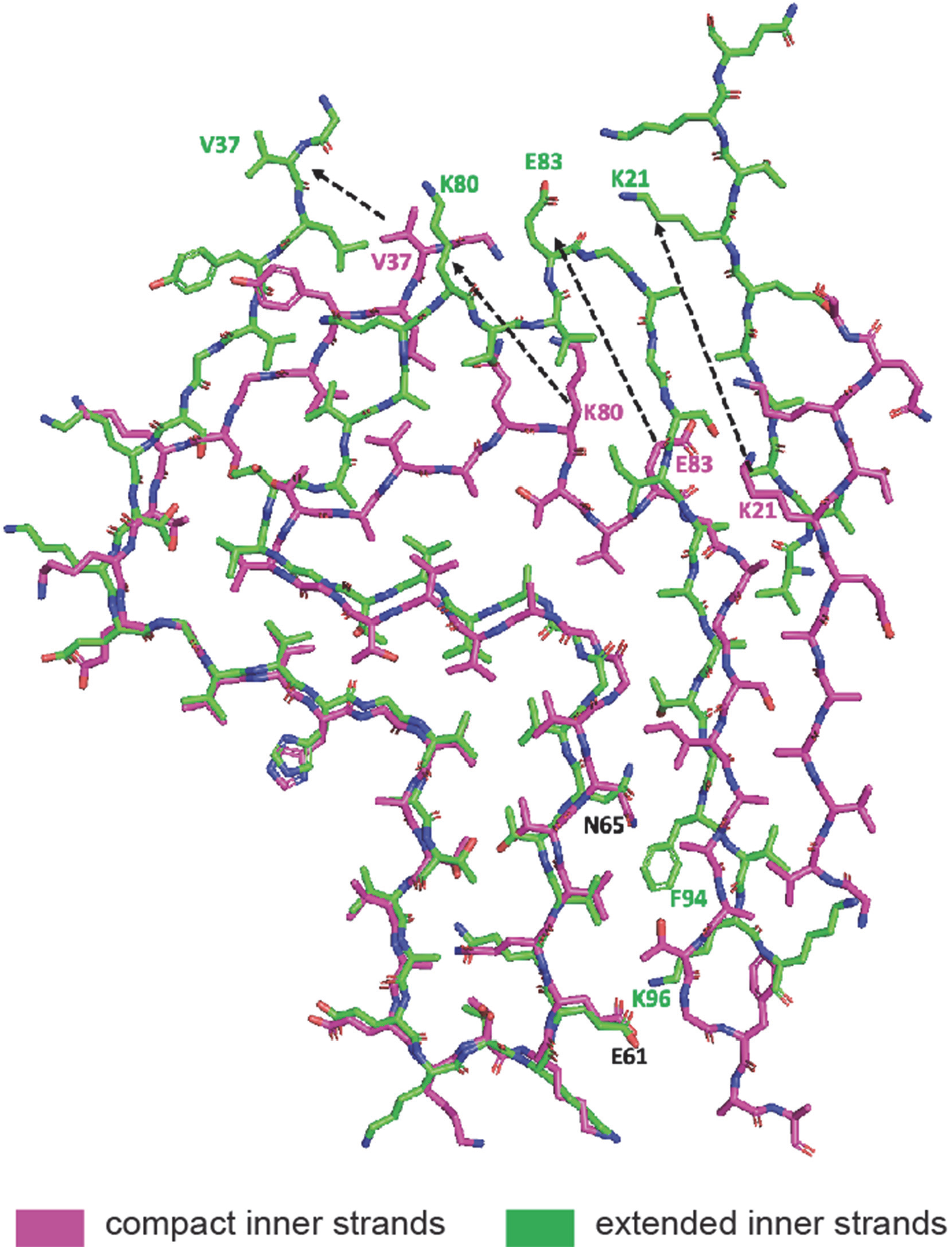
Fitted atomic model of compact and extended in both Type I and Type II β-sheet stacked filaments with variations in bonding caused by different arrangements.

**Supplemental Figure 4.**
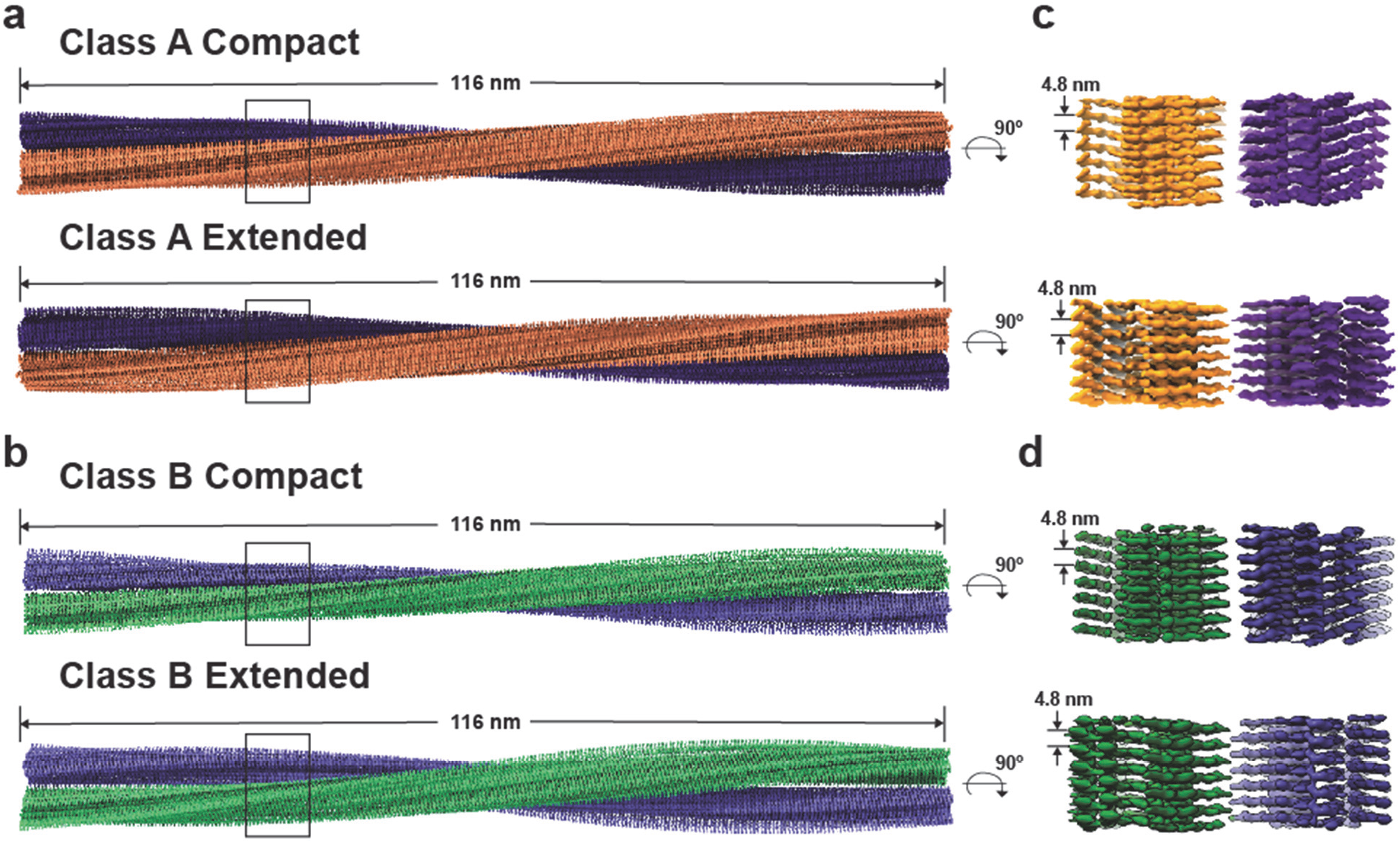
Overview of cryo-EM structures of α-synuclein fibril classes. **a.b.**, Side views of cryo-EM reconstructions for Class A and B fibrils, with extended and compact fibril arrangements. Cryo-EM density maps for both classes indicate left-handed helices with a pitch of 116 nm. Above, two protofibrils are colored dark-purple and orange for class A and light-purple and green for class B. Also shown, **c., d.,** close-up side views of the density maps of extended and compact arrangements for class A and class B fibrils with intervals indicated between each protofibril stack (common 4.8 nm).

**Supplemental Figure 5.**
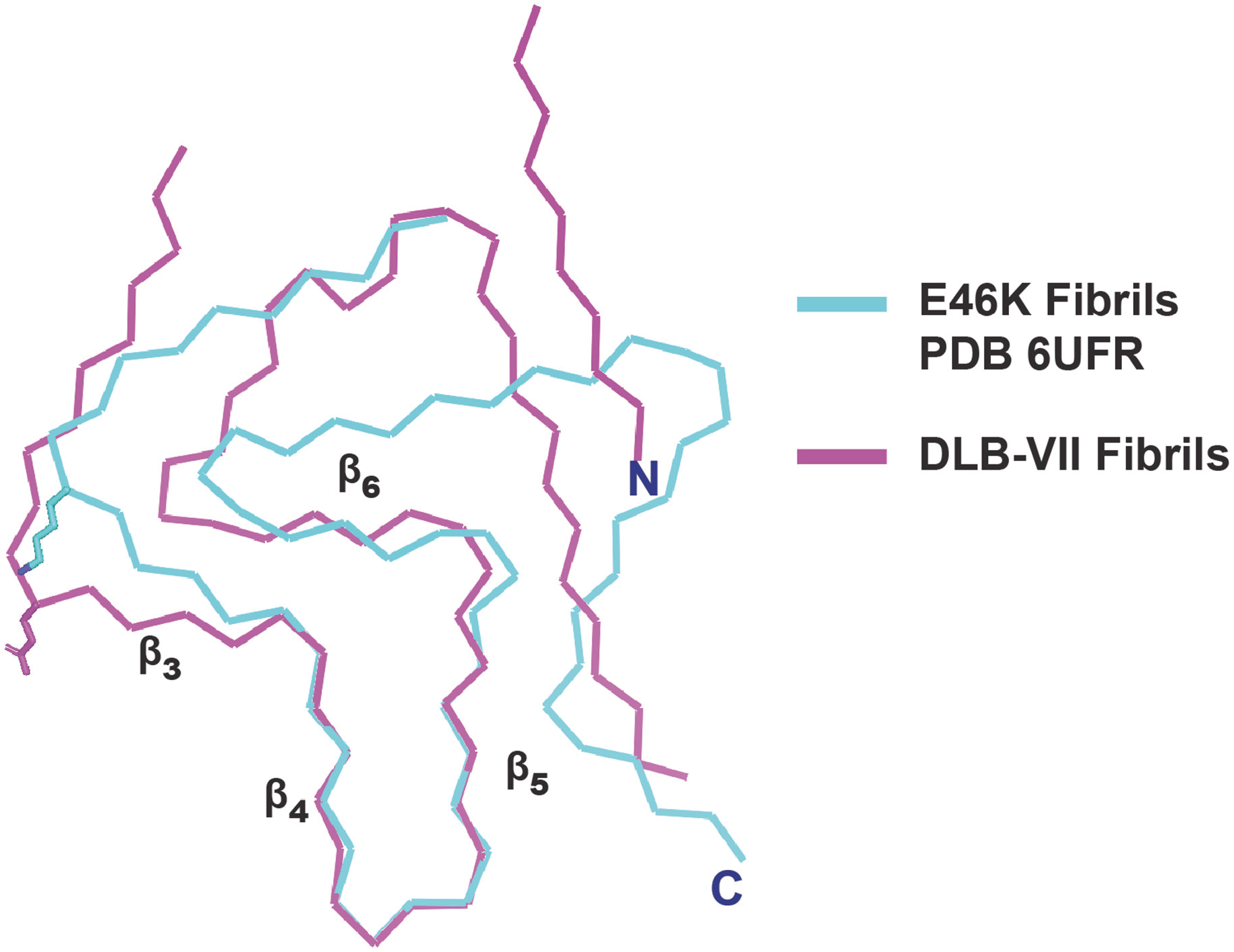
Close alignment of DLB-VII fibrils with E46K-mutated fibrils (PDB 6UFR). The mutated E46K residue is highlighted above (cyan), together with similarity in alignment from β-strands 3 to 6, centered on positive alignment between β-strands 4 and 5, with extended pocket Class B filaments from DLB-VII (purple).

**Supplemental Figure 6.**
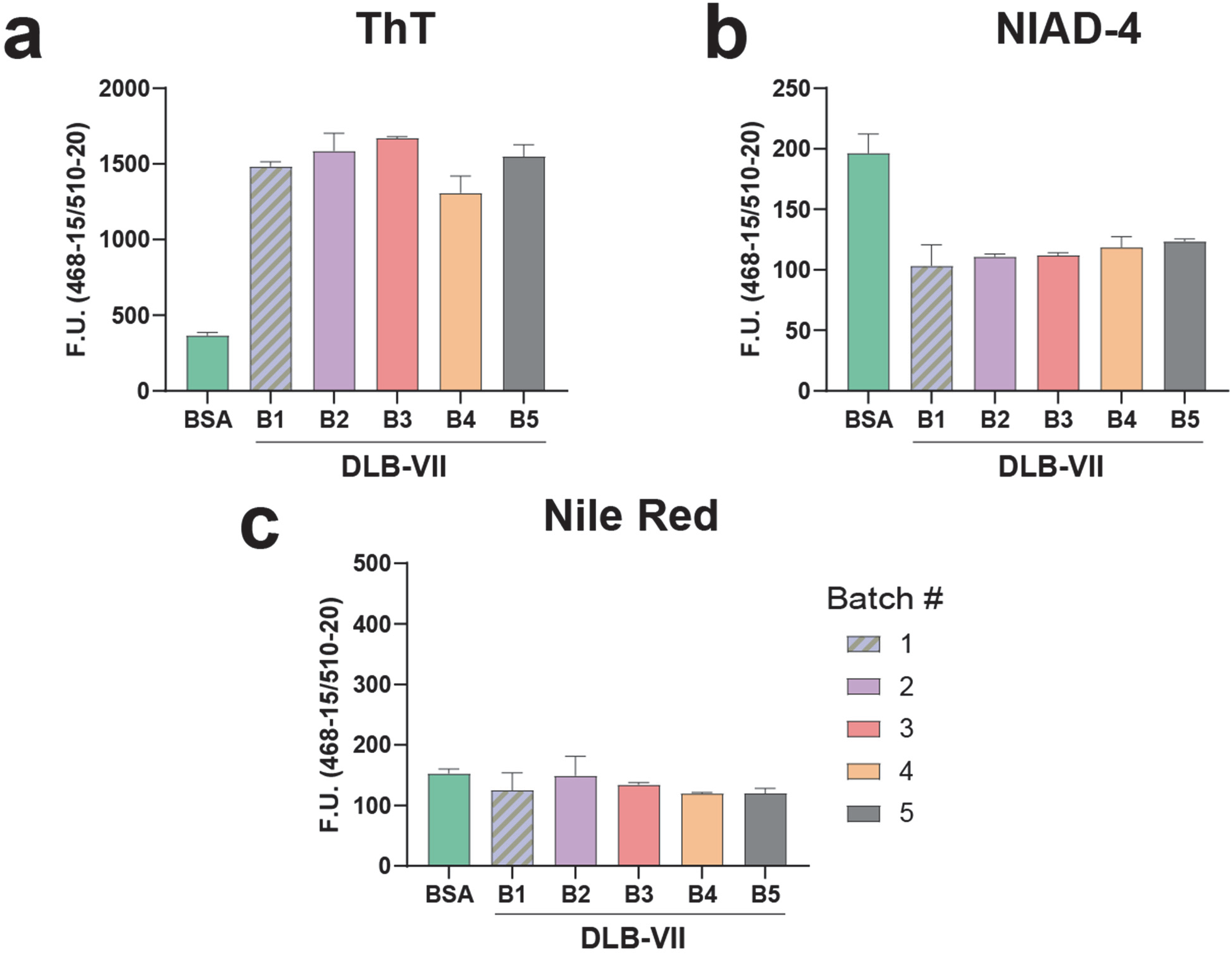
Reproducibility of dye-binding characteristics of amplified α-synuclein fibril products. **a.** Fluorescence intensity (F.U.) of five different batches (b1-5) of DLB-VII CSF (10% w/v) grown at different times and standard BSA preparation in presence of ThT, **b.** NIAD-4, or **c.** Nile Red. Columns show N=3 technical replicates from the same fibril preparation, means shown with error bars as S.E.M., One way ANOVA tests did not reveal batch differences within the spontaneous fibril group or within the DLB-VIII fibril amplification group.

**Supplemental Figure 7.**
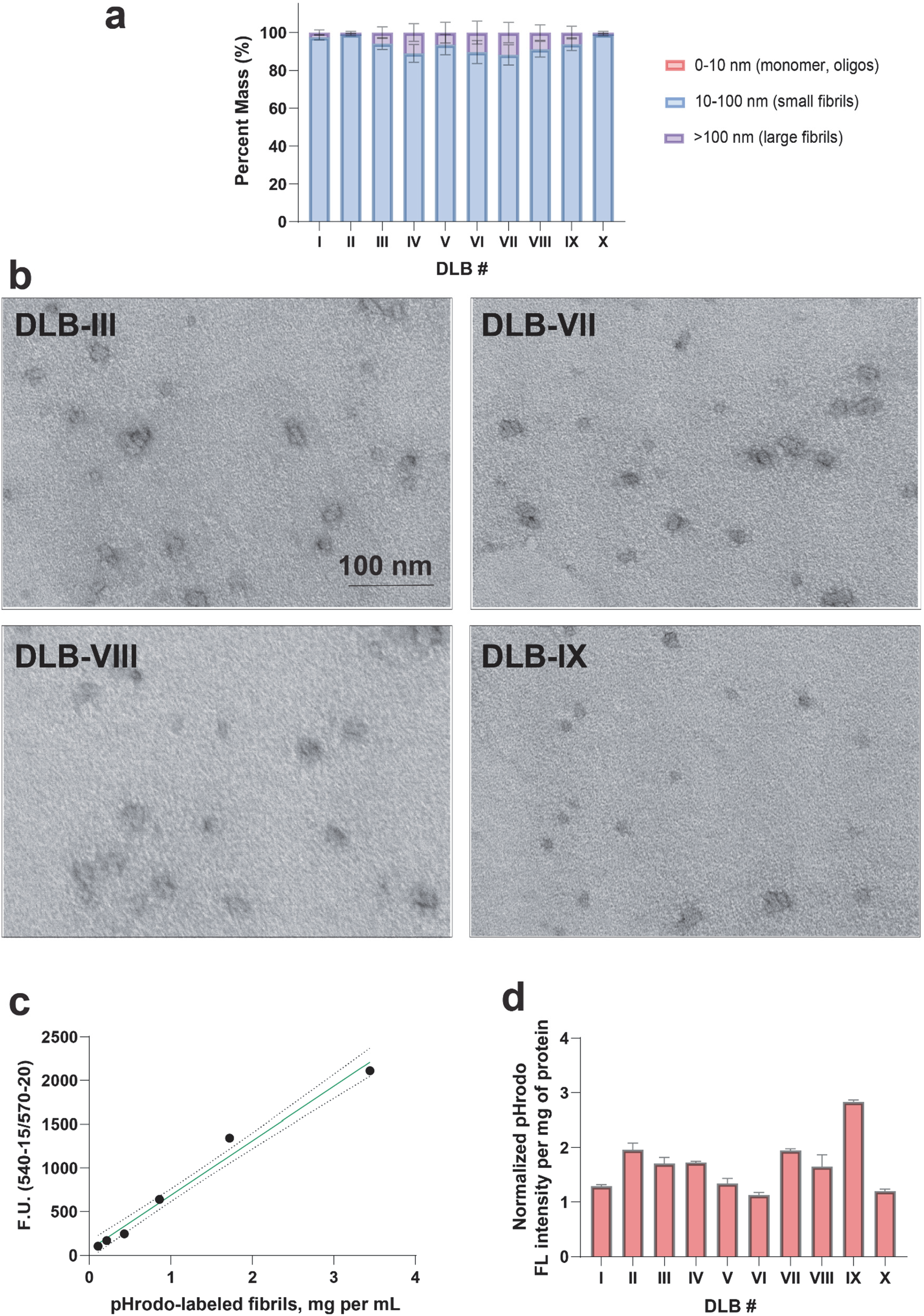
Size distributions among shredded DLB-CSF amplified α-synuclein fibrils and fluorescence intensity of pHrodo-labeled strains. **a.** Group analysis of sized populations after water-cooled sonication, with bins corresponding to 0-10 nm (e.g., monomer or small oligomeric α-synuclein), 10-100 nm (e.g., small α-synuclein fibrils), or particles larger than >100 nm (e.g., longer or aggregated fibrils). One-way ANOVA analysis did not reveal any differences across the group in triplicate measures. **b.** Selected representative negative stain electron micrographs of the processed small fibril samples prepared prior to addition to cells. Scale bar is 100 nm. **c.** Fluorescence intensity at 565 nm of pHrodo-labeled fibril samples incubated at pH 4.0 to maximize pHrodo fluorescence emission. Intensities are normalized against concentration estimated by A-280 absorbance after guanidine denaturation (see Methods-“Preparation of α-synuclein fibrils amplified from CSF”). **d.** Preceding all cellular assays, pHrodo fibril preparations were standardized between preparations to achieve similar fluorescence for each mg of fibril product. Results from three preparations are indicated as mean fluorescent values with S.E.M. as error bars.

**Supplemental Figure 8.**
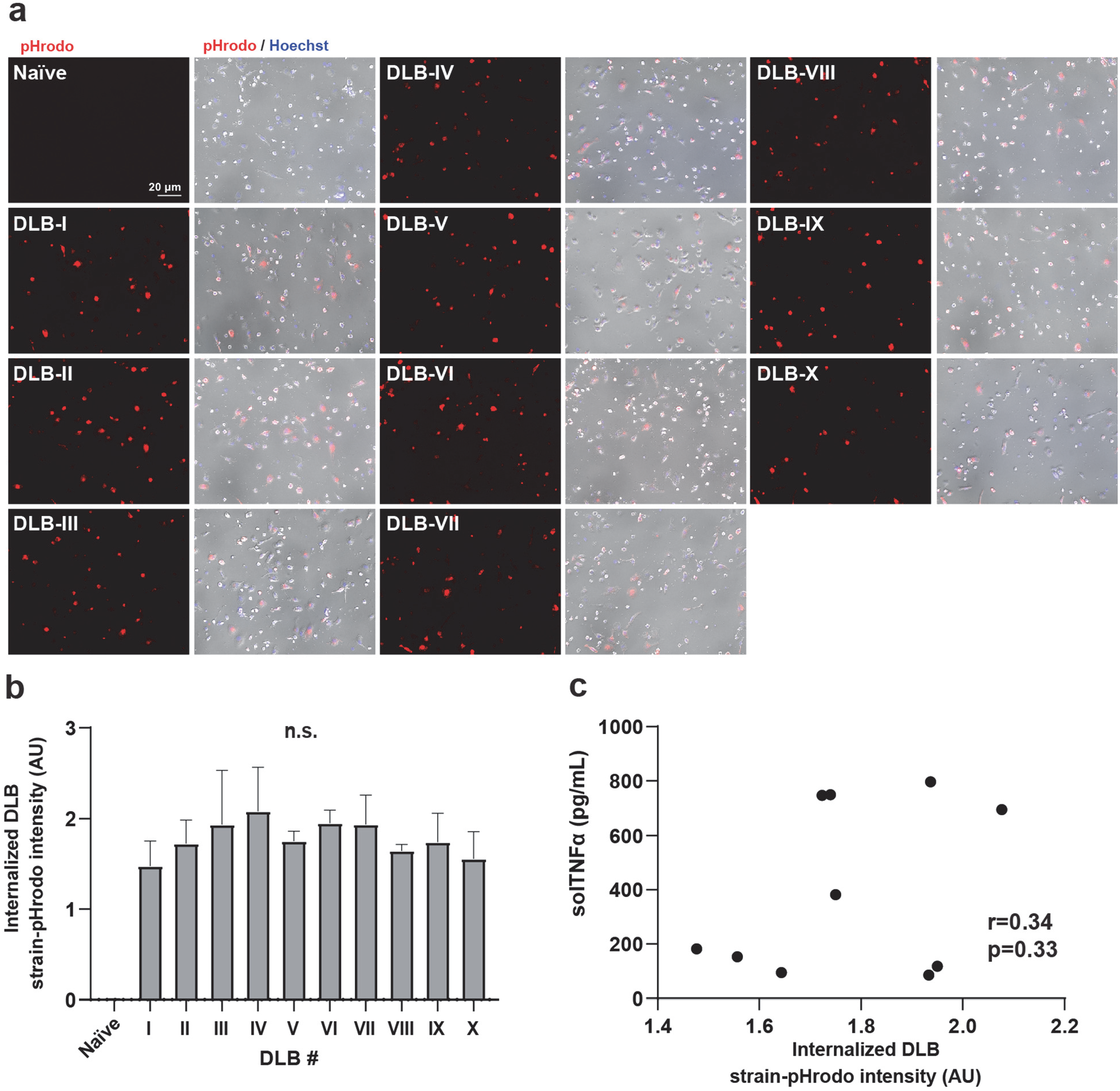
Internalization of different α-synuclein DLB-CSF amplified fibril products in primary microglia. Amplified fibril products were labeled with the pH-sensitive dye pHrodo and applied to microglia (cultured for ten days) for a period of six hours prior to widefield immunofluorescence in the living cells to resolve the amount of fibrils internalized into the endolysosomal system of the cells. **a.** Representative immunofluorescence images (red, scale bar is 20 um) with phase-contrast overlays (grey) and **b.** group analysis of fibril products. One-way ANOVA of differences between experimental conditions (N=3 independent wells per fibril treatment) did not reveal differences between DLB-I through DLB-X groups. **c.** Correlation analysis where each dot represents the mean value for fibril uptake (panel b) and stimulated TNF (**Fig. 6**).

**Supplemental Figure 9.**
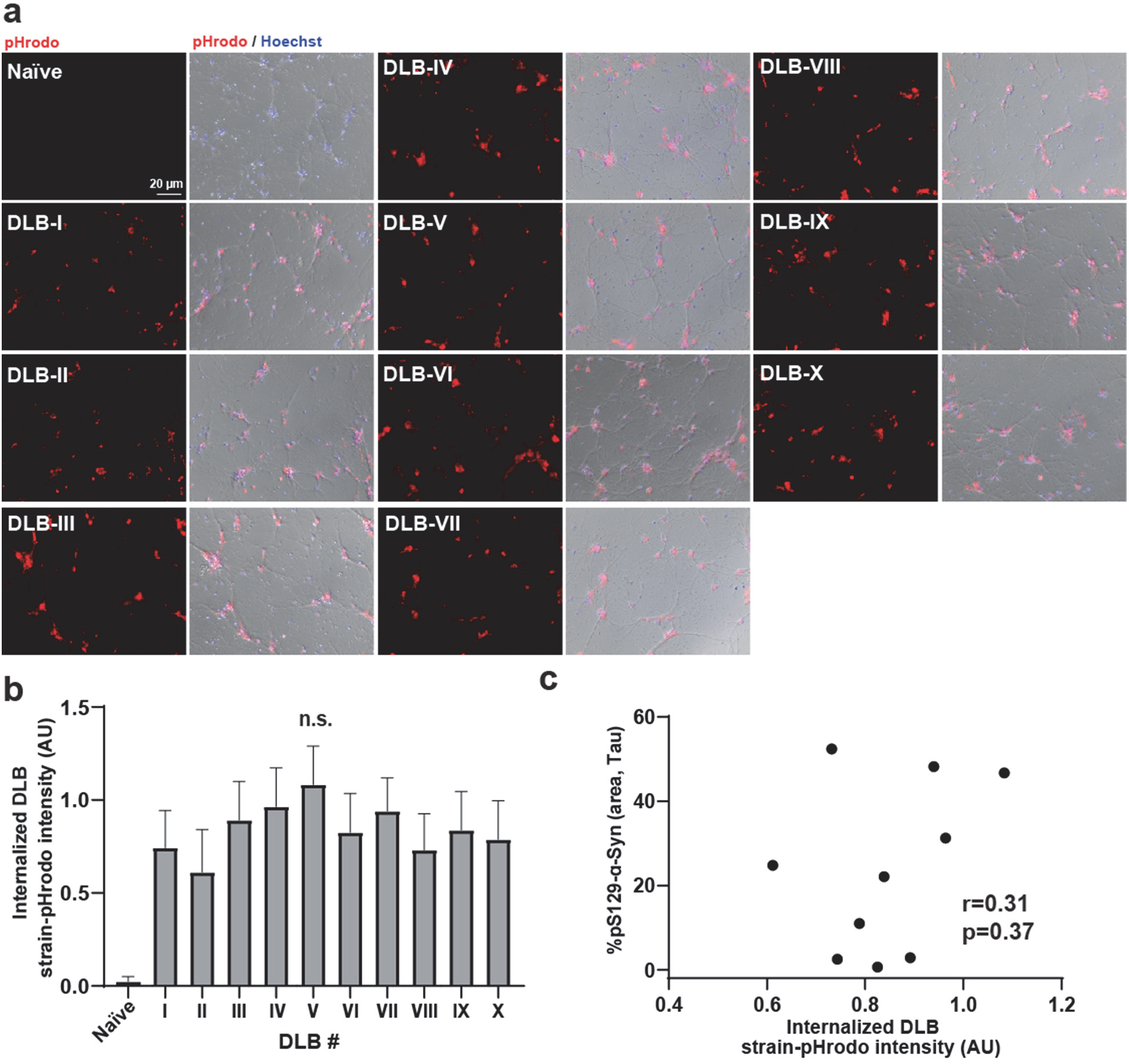
Internalization of different α-synuclein DLB-CSF amplified fibril products in primary cultured neurons. Amplified fibril products were labeled with the pH-sensitive dye pHrodo and applied to neurons (cultured for seven days) for a period of six hours prior to widefield immunofluorescence in the living cells to resolve the amount of fibrils internalized into the endolysosomal system of the neurons. **a.** Representative immunofluorescence images (red, scale bar is 20 um) with phase-contrast overlays (grey) and **b.** group analysis of fibril products. One-way ANOVA of differences between experimental conditions (N=3 independent wells per fibril treatment) did not reveal differences between DLB-I through DLB-X groups. **c.** Correlation analysis where each dot represents the mean value for fibril uptake (panel b) and potency in pS129-α-synuclein inclusion seeding (see **Fig. 7**).

**Supplemental Figure 10.**
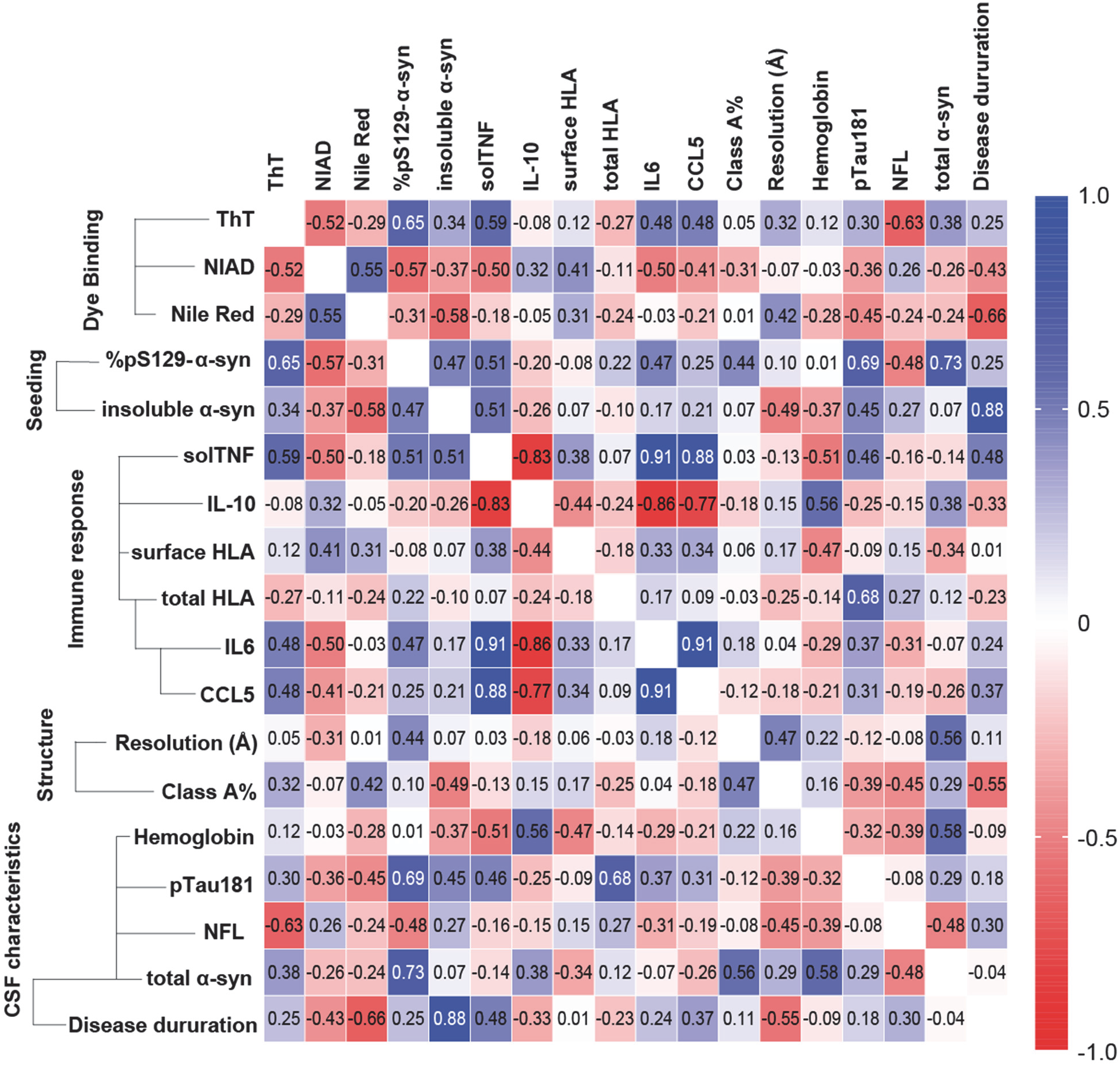
Overall correlation matrix variables associated with DLB-I through DLB-X amplified α-synuclein fibrils. Pearson’s correlation coefficients are given with darker colors showing stronger correlations, with dark red showing negative correlation and dark blue showing positive correlations.

**Supplemental Figure 11.**
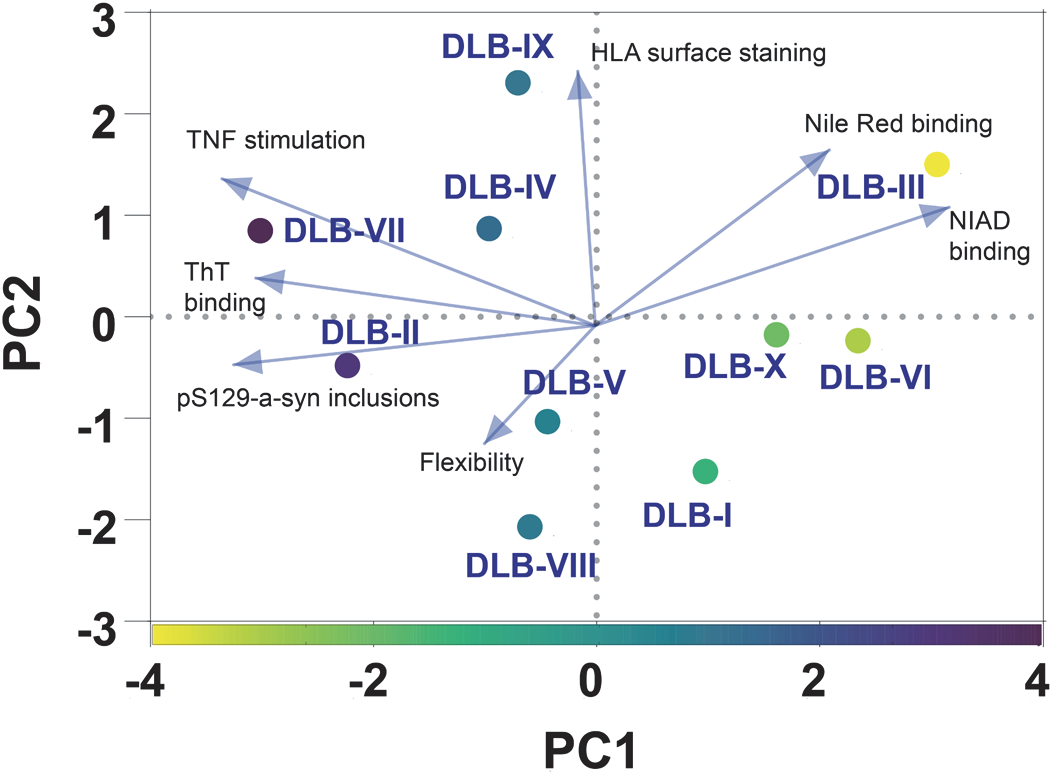
Principal component analysis (PCA) of structural and functional variabilities observed with amplified α-synuclein fibrils. PC1 and 2 show 42% and 21% proportion of variance, respectively. PC1 variances are color coded above, with variables that include dye binding profiles, seeding capacity in neurons, and immunogenic potential in microglia.

## References

1. Fanning, S., Selkoe, D. & Dettmer, U. Vesicle trafficking and lipid metabolism in synucleinopathy. Acta Neuropathol. 141, 491–510 (2021).

2. Bertoncini, C. W. et al. Release of long-range tertiary interactions potentiates aggregation of natively unstructured alpha-synuclein. Proc. Natl. Acad. Sci. U. S. A. 102, 1430–1435 (2005).

3. Braun, A. R., Lacy, M. M., Ducas, V. C., Rhoades, E. & Sachs, J. N. α-Synuclein’s Uniquely Long Amphipathic Helix Enhances its Membrane Binding and Remodeling Capacity. J. Membr. Biol. 250, 183–193 (2017).

4. Spillantini, M. G. et al. Alpha-synuclein in Lewy bodies. Nature 388, 839–840 (1997).

5. O’Nuallain, B., Williams, A. D., Westermark, P. & Wetzel, R. Seeding specificity in amyloid growth induced by heterologous fibrils. J. Biol. Chem. 279, 17490–17499 (2004).

6. Petkova, A. T. et al. Self-propagating, molecular-level polymorphism in Alzheimer’s beta-amyloid fibrils. Science 307, 262–265 (2005).

7. Jones, E. M. & Surewicz, W. K. Fibril conformation as the basis of species-and strain-dependent seeding specificity of mammalian prion amyloids. Cell 121, 63–72 (2005).

8. Luk, K. C. et al. Exogenous alpha-synuclein fibrils seed the formation of Lewy body-like intracellular inclusions in cultured cells. Proc. Natl. Acad. Sci. U. S. A. 106, 20051–20056 (2009).

9. Volpicelli-Daley, L. A. et al. Exogenous α-synuclein fibrils induce Lewy body pathology leading to synaptic dysfunction and neuron death. Neuron 72, 57–71 (2011).

10. Grozdanov, V. et al. Increased Immune Activation by Pathologic α-Synuclein in Parkinson’s Disease. Ann. Neurol. 86, 593–606 (2019).

11. Frieg, B. et al. α-Synuclein polymorphism determines oligodendroglial dysfunction. bioRxiv 2021.07.09.451731 (2021) doi:10.1101/2021.07.09.451731.

12. Long, H. et al. Wild-type α-synuclein inherits the structure and exacerbated neuropathology of E46K mutant fibril strain by cross-seeding. Proc. Natl. Acad. Sci. U. S. A. 118, (2021).

13. Marotta, N. P. et al. Alpha-synuclein from patient Lewy bodies exhibits distinct pathological activity that can be propagated in vitro. Acta Neuropathol Commun 9, 188 (2021).

14. Heerde, T. et al. Cryo-EM demonstrates the in vitro proliferation of an ex vivo amyloid fibril morphology by seeding. Nat. Commun. 13, 85 (2022).

15. Dzwolak, W., Smirnovas, V., Jansen, R. & Winter, R. Insulin forms amyloid in a strain-dependent manner: an FT-IR spectroscopic study. Protein Sci. 13, 1927–1932 (2004).

16. Cloe, A. L., Orgel, J. P. R. O., Sachleben, J. R., Tycko, R. & Meredith, S. C. The Japanese mutant Aβ (ΔE22-Aβ(1-39)) forms fibrils instantaneously, with low-thioflavin T fluorescence: seeding of wild-type Aβ(1-40) into atypical fibrils by ΔE22-Aβ(1-39). Biochemistry 50, 2026–2039 (2011).

17. Xu, H. et al. In vitro amplification of pathogenic tau conserves disease-specific bioactive characteristics. Acta Neuropathol. 141, 193–215 (2021).

18. Strohäker, T. et al. Structural heterogeneity of α-synuclein fibrils amplified from patient brain extracts. Nat. Commun. 10, 5535 (2019).

19. Peng, C. et al. Cellular milieu imparts distinct pathological α-synuclein strains in α-synucleinopathies. Nature 557, 558–563 (2018).

20. Prusiner, S. B. et al. Evidence for α-synuclein prions causing multiple system atrophy in humans with parkinsonism. Proc. Natl. Acad. Sci. U. S. A. 112, E5308–17 (2015).

21. Bargar, C. et al. Streamlined alpha-synuclein RT-QuIC assay for various biospecimens in Parkinson’s disease and dementia with Lewy bodies. Acta Neuropathologica Communications vol. 9 (2021).

22. Han, J.-Y., Jang, H.-S., Green, A. J. E. & Choi, Y. P. RT-QuIC-based detection of alpha-synuclein seeding activity in brains of dementia with Lewy Body patients and of a transgenic mouse model of synucleinopathy. Prion vol. 14 88–94 (2020).

23. Sokratian, A. et al. Heterogeneity in α-synuclein fibril activity correlates to disease phenotypes in Lewy body dementia. Acta Neuropathol. 141, 547–564 (2021).

24. Shahnawaz, M. et al. Development of a Biochemical Diagnosis of Parkinson Disease by Detection of α-Synuclein Misfolded Aggregates in Cerebrospinal Fluid. JAMA Neurol. 74, 163–172 (2017).

25. Shahnawaz, M. et al. Discriminating α-synuclein strains in Parkinson’s disease and multiple system atrophy. Nature 578, 273–277 (2020).

26. Rossi, M. et al. Ultrasensitive RT-QuIC assay with high sensitivity and specificity for Lewy body-associated synucleinopathies. Acta Neuropathol. 140, 49–62 (2020).

27. Lövestam, S. et al. Seeded assembly in vitro does not replicate the structures of α-synuclein filaments from multiple system atrophy. FEBS Open Bio 11, 999–1013 (2021).

28. Russo, M. J. et al. High diagnostic performance of independent alpha-synuclein seed amplification assays for detection of early Parkinson’s disease. Acta Neuropathol Commun 9, 179 (2021).

29. Guerrero-Ferreira, R. et al. Two new polymorphic structures of human full-length alpha-synuclein fibrils solved by cryo-electron microscopy. Elife 8, (2019).

30. Boyer, D. R. et al. The α-synuclein hereditary mutation E46K unlocks a more stable, pathogenic fibril structure. Proc. Natl. Acad. Sci. U. S. A. 117, 3592–3602 (2020).

31. Zarranz, J. J. et al. The new mutation, E46K, of alpha-synuclein causes Parkinson and Lewy body dementia. Ann. Neurol. 55, 164–173 (2004).

32. Brandenburg, E., von Berlepsch, H. & Koksch, B. Specific in situ discrimination of amyloid fibrils versus α-helical fibres by the fluorophore NIAD-4. Mol. Biosyst. 8, 557–564 (2012).

33. Harms, A. S. et al. α-Synuclein fibrils recruit peripheral immune cells in the rat brain prior to neurodegeneration. Acta Neuropathol Commun 5, 85 (2017).

34. Morales, I., Jiménez, J. M., Mancilla, M. & Maccioni, R. B. Tau oligomers and fibrils induce activation of microglial cells. J. Alzheimers. Dis. 37, 849–856 (2013).

35. Jana, M., Palencia, C. A. & Pahan, K. Fibrillar Amyloid-β Peptides Activate Microglia via TLR2: Implications for Alzheimer’s Disease. The Journal of Immunology vol. 181 7254–7262 (2008).

36. Imamura, K. et al. Distribution of major histocompatibility complex class II-positive microglia and cytokine profile of Parkinson’s disease brains. Acta Neuropathol. 106, 518–526 (2003).

37. Ryan, K. J. et al. A human microglia-like cellular model for assessing the effects of neurodegenerative disease gene variants. Sci. Transl. Med. 9, (2017).

38. Xu, E. et al. Pathological α-synuclein recruits LRRK2 expressing pro-inflammatory monocytes to the brain. Mol. Neurodegener. 17, 7 (2022).

39. Mazzulli, J. R. et al. Activation of-Glucocerebrosidase Reduces Pathological-Synuclein and Restores Lysosomal Function in Parkinson’s Patient Midbrain Neurons. Journal of Neuroscience vol. 36 7693–7706 (2016).

40. Abdelmotilib, H. et al. α-Synuclein fibril-induced inclusion spread in rats and mice correlates with dopaminergic Neurodegeneration. Neurobiol. Dis. 105, 84–98 (2017).

41. Luk, K. C. et al. Molecular and Biological Compatibility with Host Alpha-Synuclein Influences Fibril Pathogenicity. Cell Rep. 16, 3373–3387 (2016).

42. Kuo, Y.-M. et al. Extensive enteric nervous system abnormalities in mice transgenic for artificial chromosomes containing Parkinson disease-associated alpha-synuclein gene mutations precede central nervous system changes. Hum. Mol. Genet. 19, 1633–1650 (2010).

43. Fenyi, A. et al. Seeding Propensity and Characteristics of Pathogenic αSyn Assemblies in Formalin-Fixed Human Tissue from the Enteric Nervous System, Olfactory Bulb, and Brainstem in Cases Staged for Parkinson’s Disease. Cells 10, (2021).

44. Schweighauser, M. et al. Structures of α-synuclein filaments from multiple system atrophy. Nature 585, 464–469 (2020).

45. Chandra, R., Hiniker, A., Kuo, Y.-M., Nussbaum, R. L. & Liddle, R. A. α-Synuclein in gut endocrine cells and its implications for Parkinson’s disease. JCI Insight 2, (2017).

46. Scheres, S. H. W. RELION: implementation of a Bayesian approach to cryo-EM structure determination. J. Struct. Biol. 180, 519–530 (2012).

47. Rohou, A. & Grigorieff, N. CTFFIND4: Fast and accurate defocus estimation from electron micrographs. J. Struct. Biol. 192, 216–221 (2015).

48. Wagner, T. et al. SPHIRE-crYOLO is a fast and accurate fully automated particle picker for cryo-EM. Commun Biol 2, 218 (2019).

49. Emsley, P., Lohkamp, B., Scott, W. G. & Cowtan, K. Features and development of Coot. Acta Crystallogr. D Biol. Crystallogr. 66, 486–501 (2010).

50. Liebschner, D. et al. Macromolecular structure determination using X-rays, neutrons and electrons: recent developments in Phenix. Acta Crystallographica Section D: Structural Biology 75, 861–877 (2019).

51. Davis, I. W. et al. MolProbity: all-atom contacts and structure validation for proteins and nucleic acids. Nucleic Acids Res. 35, W375–W383 (2007).

52. Friesner, R. A. et al. Extra precision glide: docking and scoring incorporating a model of hydrophobic enclosure for protein-ligand complexes. J. Med. Chem. 49, 6177–6196 (2006).

53. Halgren, T. A. et al. Glide: a new approach for rapid, accurate docking and scoring. 2. Enrichment factors in database screening. J. Med. Chem. 47, 1750–1759 (2004).

54. Friesner, R. A. et al. Glide: a new approach for rapid, accurate docking and scoring. 1. Method and assessment of docking accuracy. J. Med. Chem. 47, 1739–1749 (2004).

